# Stromal cell sialylation suppresses T cells in inflammatory tumour microenvironments: A new tumour stromal cell immune checkpoint?

**DOI:** 10.1101/2021.06.18.447879

**Authors:** Hannah Egan, Oliver Treacy, Kevin Lynch, Niamh A Leonard, Grace O’Malley, Kim De Veirman, Karin Vanderkerken, Michael Craughwell, Laurence J Egan, Thomas Ritter, Aisling M Hogan, Keara Redmond, Margaret Sheehan, Aoife Canney, Sean Hynes, Emma Kerr, Philip D Dunne, Michael E O’Dwyer, Aideen E Ryan

## Abstract

Immunosuppressive tumour microenvironments (TME) reduce the effectiveness of immune responses in cancer. Non-haematopoietic mesenchymal stromal cells (MSC), the precursor to cancer associated fibroblasts (CAFs), dictate tumour progression by enhancing immune cell suppression. Hyper-sialylation of glycans promotes immune evasion in cancer, but the role of sialyation in stromal cell-mediated immunosuppression is unknown.

Here we study changes in sialyltransferase (ST) enzymes and associated surface expressed sialic acid in stromal cells following inflammatory and tumour secretome conditioning. We show that tumour conditioned stromal cells have increased levels of sialyltransferases, α2,3/6 linked sialic acid and siglec ligands. In tumour models of solid (colorectal cancer) and haematological (multiple myeloma) stromal rich tumours, stromal cell sialylation is associated with enhanced immunosuppression. Using datasets and patient samples, we confirm that targeting sialylation in tumour stromal cells reverses immune cell exhaustion. Targeting stromal cell sialylation may represent a novel immune checkpoint to reactivate anti-tumour immunity.

## Introduction

Immunosuppressive tumour microenvironments (TME) reduce the effectiveness of immune based therapies for the treatment of cancer [1]. Colorectal cancer can be classified into subtypes based on distinct molecular and clinical features [2]. In colorectal cancer, the heterogeneous environment is composed of numerous cellular components that can facilitate an immunosuppressive microenvironment, including immune, stromal, and endothelial cells. In colon cancer, the type, density, and location of immune cells within tumours can predict clinical outcome and tumour recurrence [3]. Lower densities of CD3, CD8 immune cells identify patients with tumour recurrence compared to patients whose tumours did not recur [3, 4]. More recently, a mesenchymal signature in CRC, reflecting an increased stromal cell content compared with other CRC subgroups is associated with poor prognosis [5, 6]. Similarly, studies of multiple myeloma, a cancer that arises in the bone marrow in a stromal cell dense environment, indicate the complex role of stromal cells in regulating tumour progression and immune evasion [7]. These observations indicate the importance of understanding the key cellular and molecular events that dictate immunosuppressive tumour microenvironments and tumour extrinsic mechanisms that influence tumour progression.

Non-haematopoietic intestinal mesenchymal stromal cells (MSC), the precursor to cancer associated fibroblasts (CAFs) are major components of the CRC and MM TME [8]. They are a heterogeneous population of stromal cells of mesenchymal lineage, defined by a combination of morphological characteristics, tissue origin and lack of lineage markers [9, 10]. Stromal cells in the tumour microenvironment arise from local tissue mesenchymal stromal cells, fibroblasts, trans-differentiation events and recruited bone marrow MSC and [11]. The definition of cell types and subtypes within the classification of stromal cells is complicated by the lack of distinct markers [9]. Both the high proportion, localization and function of MSC in CRC, and other stromal rich tumours suggest that these cells are crucial to tumor development [10]. MSC are positioned between the epithelial cells and the underlying vasculature and can passively or actively impair immune cell trafficking and activation [8, 12]. The immunological hallmarks of stromal cells in the TME include regulation of immune cell infiltration, regulation of anti-tumour immune responses and responsiveness to immunotherapy [9]. Stromal cell signatures in colon tumours and MM are associated with tumour progression and a poorer prognosis, immune evasion and therapy resistance [6, 13, 14]. It is unclear whether this association is due to inherent tumour promoting or immunosuppressive functions in the TME [8]. This knowledge highlights the need to investigate mechanisms to improve discovery of effective stroma-targeting therapeutic strategies.

Tumour promoting inflammation, an acknowledged hallmark of cancer is associated with tumour initiation and progression. We, and others, have shown that inflammatory signalling initiated by TNF-α in CRC promotes stromal cell mediated immunosuppression and CRC progression in vivo [15]. Recent data in MM links inflammatory stromal cell landscapes with MM survival and immune modulation [16]. Studies have shown that targeting PD-L1, PD-L2, FasL and PGE2 can reverse stromal mediated immunosuppression in the TME; however identifying the predominant immunosuppressive mechanisms within varied TMEs remains a challenge [17]. Chronic inflammation can alter glycosylation and emerging knowledge on the role of glycosylation in tumour progression, indicates its association with poorer prognosis [18, 19]. One of the more common changes in cancer glycosylation is an upregulation of sialylated glycans termed hyper-sialylation [20]. Sialic acid is a common component of glycan molecules, and its presence can result in altered protein function and immune recognition[20]. Cancer cells upregulate sialic acid expression, which they can hijack to evade immune clearance [21, 22]. However, little is known about the sialylation profiles of stromal cells and the functional consequences of these on immune cells in inflammatory TMEs.

Sialo glycans are recognised by the Siglec receptors (Sialic acid binding immunoglobulin-type lectins) which are expressed on the surface of both innate and adaptive immune cells [23]. Siglecs are a class of self-pattern recognition receptors (SPPRs) that co-regulate the function of immune cells. Inhibitory Siglecs contain tyrosine-based inhibitory signalling motifs (ITIMs) that mediate inhibitory signals upon binding sialogycans [23, 24]. Hyper-sialylation of glycans is linked to increased immune evasion, drug resistance, tumour invasiveness and metastasis [20, 25]. The inhibitory Siglecs most strongly implicated in immune evasion in cancer are Siglec-7, Siglec-9 and Siglec-10, which are expressed on natural killer (NK) cells, macrophages, and T cells, respectively [25-28]. Recent conflicting reports regarding the pro and anti-tumour effects of targeting Siglec ligands, specifically on tumour cells have indicated the complexity of sialic acid dependent signalling in the regulation of tumour growth [29, 30]. This data reflects the heterogeneity of cell type specific sialyation in the TME. Overcoming immunosuppression and enhancing immunotherapy responses is a key challenge in cancer treatment.

We propose that sialylation of the stromal compartment of the TME is an unexplored immunological target to reverse TME immunosuppression.We hypothesised that inflammation and the inflammatory TME induces stromal cell sialylation, which in turn regulates stromal cell mediated immunosuppression. Here we show, using two preclinical tumor models and two clinical patient cohorts, that stromal cell sialylation can suppress T cell activation, function and phenotype through cell-cell contact dependent mechanisms. We show that inflammation in the TME can enhance stromal cell sialylation, siglec 7 and 9 ligand expression, which is associated with T cell suppression and exhaustion. Targeting sialyl-transferase activity in stromal cells inhibits siglec ligand expression, reverses T cell suppression and exhaustion, and represents a novel immunological target in stromal dense tumors. Targeting stromal cell sialylation and/or siglec/siglec ligand interactions, in combination with other immune cell-based immunotherapies represents an innovative strategy to enhance anti-tumor immunity in immunosuppressive TMEs.

## Materials and Methods

### Mesenchymal stromal cell isolation and culture

Balb/c or C57BL/6 mice were purchased from Envigo Laboratories (Oxon, UK) or Jackson Laboratories (Bar Harbor, Maine) and housed and maintained following the conditions approved by the Animals Care Research Ethics Committee of the National University of Ireland, Galway (NUIG) and conducted under individual and project authorisation licenses from the Health Products Regulatory Authority (HPRA) of Ireland. All animals were housed and cared for under Standard Operating Procedures of the Animal Facility at the Biomedical Sciences Biological Resource Unit, NUIG. For murine mesenchymal stromal cell (MSC) isolation, Balb/c or C57BL/6 mice were euthanized by CO_2_. The femur and tibia were removed, cleaned of connective tissue and MSC were flushed from the bones. Bone marrow derived cells were filtered and plated at a density of 1×10^6^ cells per T175 flask (Sarstedt, Wexford, Ireland). Cells were incubated at 37°C in normoxia (21% O_2_) for Balb/c MSC or hypoxia (5% O_2_) for C57BL/6 MSC in culture medium consisting of MEM-α (ThermoFisher Scientific, Dublin, Ireland) supplemented with 10% heat-inactivated foetal bovine serum (FBS) (Sigma-Aldrich, Wicklow, Ireland), and 1% penicillin/streptomycin (ThermoFisher Scientific). Non-adherent cells were removed 24 hours later and cells were refreshed with new media. This process was repeated until cells reached confluency. MSC were characterized according to the criteria set out by the International Society for Cellular Therapy (ISCT). Cell surface characterization and tri-lineage differentiation was performed [15, 31].

### Mouse Cell lines

CT26 mouse colon adenocarcinoma cells, derived from Balb/c mice, were purchased from the European Collection of Cell Cultures (ECACC) and cultured in DMEM media (ThermoFisher Scientific) supplemented with 10% FBS (Sigma-Aldrich) and 1% penicillin/streptomycin (ThermoFisher Scientific). The MC38 murine colon adenocarcinoma cell line, derived from C57BL/6 mice, was purchased from Kerafast (MA, USA) and cultured in DMEM medium (Sigma) supplemented with 10% FBS (Sigma-Aldrich), 1% penicillin/streptomycin (ThermoFisher Scientific), 2mM glutamine, 0.1mM nonessential amino acids, 1mM sodium pyruvate, 10mM Hepes and 50ug/ml gentamycin (all ThermoFisher Scientific).

### Generation of tumour cell secretome, MSC-conditioning and sialyl-transferase inhibition

To generate inflammatory-conditioned MSC, TNF-α + IL-1β (50ng/ml of each; Peprotech, London, UK) were added to MSC medium and cells were incubated at 37°C in normoxia for 72 hours. Cells were washed twice with Dulbecco’s phosphate buffered saline (DPBS) (ThermoFisher Scientific), stained for cell surface expression of target markers (See Table S1) and analyzed by flow cytometry or used in T cell co-culture assays. For generation of tumour cell secretome (TCS), CT26, MC38, HT29, HCT116, RPMI 8226 or MM1S cells were seeded in T175 flasks at a density of 1×10^6^, 8×10^5^, 2×10^6^, 1.5×10^6^, 3×10^5^ or 5×10^5^ cells/ml, respectively, in cell culture medium. Cells were grown at 37°C in normoxia for a total of 72 hours, whereupon the conditioned medium was collected and spun at 1000 x g. The pellet was discarded, and the conditioned medium was stored at -80°C. For generation of inflammatory TCS (iTCS), TNF-α (10-100ng/ml, Peprotech) was added to tumour cell cultures 24 hour prior to conditioned medium collection. For MSC conditioning, cells were seeded at a density of 0.03×10^6^ (for mouse cancer cells) or 0.06×10^6^ (for human cancer cells) per well of a 6-well plate in 2ml of appropriate MSC culture medium. 24 hours after seeding, MSC medium was removed and replaced with 40% fresh MSC medium + 60% TCS/iTCS. TCS/iTCS-conditioned MSC were collected, washed twice in DPBS (ThermoFisher Scientific), stained for cell surface expression of target markers (See Table S1) and analyzed by flow cytometry or used in T cell co-culture assays.

For experiments involving sialyl-transferase inhibition (SI), MSC were cultured for 72 hours with TNF-α + IL-1β/TCS/iTCS. MSC were counted and re-seeded at 0.03×10^6^ per well of a 6-well plate. Conditioned MSCs were plated for 4 hours before medium was removed and replaced with either fresh MSC medium supplemented with 50ng/ml each of TNF-α + IL-1β or 40% fresh MSC medium + 60% TCS/iTCS + 200uM of the SI, 3F_ax_Neu5Ac (Bio-Techne).Conditioning was repeated twice over a period of 6d. TNF-α + IL-1β/TCS/iTCS-conditioned MSC ± SI were then collected, washed twice in DPBS (ThermoFisher Scientific), stained for cell surface expression of target markers (See Table S1) and analysed by flow cytometry or used in T cell co-culture assays. For SI treatment of NAFs/CAFs, cells were seeded at 0.06×10^6^ per well of a 6-well plate in 2ml of MSC media. Cells were plated for 4 hours, followed by addition of 200uM of 3FaxNeu5Ac (Bio-Techne). Conditioning was repeated twice over a period of 6d. NAFs/CAFs ± SI were incubated for a further 72 hours before they were collected, washed twice in DPBS (ThermoFisher Scientific), stained for cell surface expression of target markers (See Table S1) and analysed by flow cytometry or used in PBMC co-culture assays.

### Cell surface characterization by flow cytometry

Prior to antibody/lectin staining, MSC and cancer cells were trysinised, counted and washed twice in FACS buffer (DPBS supplemented with 2% FBS and 0.05% sodium azide (all Sigma-Aldrich)). For cell surface characterization, 5×10^4^ MSC or cancer cells were stained for the specific markers. See Table S1 for full details of antibodies and lectins used. Samples were analysed using a BD FACSCanto II Flow Cytometer (BD Biosciences, California, USA). Flow cytometry data was analysed using FlowJo analysis software version 10 (Tree Star Inc., OR, USA).

### Stromal cell RNA Sequencing

RNA sequencing (RNA-Seq) was performed on MSC^Control^, MSC^TNFα+IL-1β^, CT26 MSC^TCS^ and CT26 MSC^iTCS^ cells according to our published protocol [31]. RNA sequencing and data analysis were outsourced to Arraystar (USA).

### Mouse lymphocyte isolation

To obtain primary Balb/c and C57BL/6 lymphocytes, lymph nodes and spleens were harvested from healthy Balb/c and C57BL/6 mice following CO_2_ euthanasia. Single cell suspensions of lymph nodes and spleens were prepared by gentle mashing of the organs through a 40μM cell strainer (ThermoFisher Scientific) in 6cm petri dishes (Sarstedt) containing 5ml DPBS (ThermoFisher Scientific). The single cell suspensions were centrifuged at 800 x g for 5 mins. Lymphocytes were washed in DPBS and counted. Splenocytes were re-suspended in erythrocyte lysis buffer (distilled water, 0.15M NH_4_CL, 10mM KHCO_3_, Sodium EDTA 0.1mM) and incubated on ice for 5 mins. The reactions were stopped by adding complete medium consisting of RPMI 1640 (ThermoFisher Scientific) supplemented with 10% heat-inactivated FBS, 1% sodium pyruvate, 1% non-essential amino acids, 1% L-glutamine, 1% penicillin/streptomycin and 0.1% β-mercaptoethanol (all Sigma-Aldrich). Cells were centrifuged at 800 x g for 5 mins, washed twice in DPBS, re-suspended in culture medium and included in immunosuppression assays.

### Mouse co-culture assays

For lymphocyte proliferation assays, lymph nodes and spleens were isolated from Balb/c mice and single cell suspensions prepared as described above. Cells were re-suspended in complete medium (RPMI 1640 (ThermoFisher Scientific) supplemented with 10% heat-inactivated FBS, 1% sodium pyruvate, 1% non-essential amino acids, 1% L-glutamine, 1% penicillin/streptomycin and 0.1% β-mercaptoethanol (all Sigma-Aldrich)). Cells were stained with the CellTrace™ Violet proliferation kit (ThermoFisher Scientific) according to the manufacturer’s protocol and seeded in 96 well round bottom plates (Sarstedt) at a concentration of 2×10^5^ cells/100μl of complete medium with or without anti-mouse anti-CD3/CD28 Dynabeads (ThermoFisher Scientific) at a ratio of 1:4 (beads: lymphocytes). TNF-α + IL-1β/TCS/iTCS or control MSC ± SI were added to wells of lymphocytes at a concentration of 2×10^4^ cells/100μl (ratio of 1:10 MSC:lymphocytes) of mouse MSC medium. After 96 hours, supernatants were collected from the co-cultures and stored at -80°C. Cells were incubated with the following. anti-mouse antibodies (see Table S1 for additional details) diluted. in FACS buffer (DPBS supplemented with 1% FBS and 0.05% sodium azide (Sigma-Aldrich)): CD3-FITC, CD4-PE/Cy7, CD8-APC/Cy7, CD25-PE, CD69-PE (all Biolegend) and Siglec E-APC (Bio-Techne). Samples were analysed using a BD FACSCanto II Flow Cytometer (BD Biosciences). Flow cytometry data was analysed using FlowJo analysis software version 10 (Tree Star Inc.).

### ELISAs and Griess Assay for NO Quantification

Supernatants from MSC and T-cell cocultures were analyzed using IL-10 Ready SET-Go! ELISA kits kits (Affymetrix, eBioscience) for secretion of IL-10 as previously described [31]. IL-6 was detected in culture supernatants from MSC/T cell co-culture assays by magnetic Luminex assays (R&D Systems, Biotechne, Abingdon, UK) as previously described [31]. PGE2 was detected using an individual PGE2 assay (R&D Systems, Biotechne) as previously described [31]. NO concentration was quantified in culture supernatants by a Griess assay. 100 mL of supernatant was combined with an equal volume of Griess reagent (composed of 1% sulfanilamide and 0.1% N-1-(naphthyl)ethylenediamine dihydrochloride in 2.5% H3PO4) (Abcam, Cambridge, UK) in a 96-well flat bottom plate [32]. Absorbance was measured at 540 nm on a plate reader (PerkinElmer, Ireland).

### 5T33MM Myeloma Mouse Model

To generate multiple myeloma derived-MSC (MM-MSC), C57BL/KaLwRij mice were purchased from Envigo Laboratories and housed and maintained following the conditions approved by the Ethical Committee for Animal Experiments, Vrije Universiteit Brussel (license no. LA1230281, 16-281-6). The 5T33MM model originated from spontaneously developed MM in elderly C57BL/KalwRij mice and was established by intravenous transfer of the diseased marrow into young syngeneic mice. For *in vivo* experiments, mice were intravenously inoculated with 5×10^5^ 5T33MM cells [33, 34]. To generate MSC, total bone marrow was isolated from naive and diseased 5T33MM mice followed by red blood cell lysis. Tumor load in the 5T33MM mice was above 80%, as determined by cytospin staining and calculation of the percentage plasmacytosis (data not shown). Cells were cultured as described above for wild-type (WT) mice. To generate myeloma conditioned medium, 5T33vt cells, clonally identical to the in vivo model, were cultured at a concentration of 10^6^ cells per mL in RPMI 1640 medium with 10% FCS for 48 hours. 5T33vt supernatant was used either concentrated 1X or concentrated 10x using a centrifugal filter (3kDa molecular weight cutoff) (Millipore). MSC were conditioned using 1X 5T33vt or 10X 5T33vt TCS in vitro.

### Human Tumour cell lines

HT29 human colorectal adenocarcinoma cells and HCT116 human colorectal carcinoma cells were purchased from the American Type Culture Collection (ATCC, VA, USA) and cultured in McCoy’s 5A medium (Sigma-Aldrich) with 10% FBS (Sigma-Aldrich), 1% L-glutamine and 1% penicillin/streptomycin (both ThermoFisher Scientific). All cell lines were expanded, frozen and used within 15 passages.

### Isolation of Human Mesenchymal Stromal cells

Human MSC (hMSC) were isolated and expanded [15]. Briefly, hMSC were isolated from the bone marrow of three healthy volunteers at Galway University Hospital under an ethically approved protocol (NUIG Research Ethics Committee, Ref: 08/May/14) according to a standardized procedure. Written consent was obtained from the volunteers. Bone marrow cell suspensions were layered onto a Ficoll density gradient. The nucleated cell fraction was collected, washed, and resuspended in MSC culture medium. 24 hours later, non-adherent cells were removed, and fresh medium was added. Individual colonies of fibroblast-like cells were allowed to expand and approach confluence prior to passage. hMSC were grown in MEM-α (ThermoFisher Scientific, Dublin, Ireland) supplemented with 10% FBS (Sigma-Aldrich), 1% penicillin/streptomycin (ThermoFisher Scientific) and fibroblast growth factor 2 (FGF2, 1ng/ml; Sigma-Aldrich).

### Isolation of primary stromal cells from colorectal cancer patient tumours and adjacent normal mucosa

Colorectal tumor and adjacent normal mucosal tissue were obtained from patients undergoing colon tumor resection at University Hospital Galway under an ethically approved protocol (Clinical Research Ethics Committee, Ref: C.A. 2074). Written informed explicit consent was obtained from all patients prior to sampling. Following pathological assessment, biopsies of central tumour and normal mucosal tissue were removed and washed intact 5 times with Hank’s balanced salt solution (HBSS) supplemented with 10% penicillin/streptomycin (both Sigma-Aldrich). After washing, the biopsies were cut into 2-3mm pieces and dissociated using a human Tumour Dissociation Kit (Miltenyi Biotec, Surrey, UK) according to the manufacturer’s protocol with some modifications. Incubation time was reduced to 2 hours and dissociation was achieved by inverting the suspensions 10 times every 30 mins. The resultant cell suspensions were filtered through 70µm cell strainers (ThermoFisher Scientific) and centrifuged at 400 x g for 5 mins. Single cell suspensions were re-suspended in complete human MSC medium (RPMI 1640 medium (ThermoFisher Scientific) supplemented with 10% heat-inactivated FBS, 1% sodium pyruvate, 1% HEPES solution, 1% L-glutamine, 1% penicillin/streptomycin, 0.1% β-mercaptoethanol and 1ng/ml FGF2 (all Sigma-Aldrich)). Cells were then seeded in 6 well plates (Sarstedt) until stromal cell colony establishment was observed. Stromal cells isolated from colorectal tumour biopsies were termed cancer-associated fibroblasts (CAFs), while stromal cells derived from patient-matched normal mucosal tissue were termed normal-associated fibroblasts (NAFs).

### Immunohistochemical analysis human colorectal cancer samples

Sections were taken from formalin fixed paraffin embedded (FFPE) tumour tissue blocks cut between 3-5 µM thick, with a rotary microtome. Sections were stained with haematoxylin and eosin. For immunohistochemistry, both CD3 (Thermofisher) and CD8 (Dako) markers were used on the automated BondMax system (Vision Biosystems). The slides were scanned using an Olympus (VS120) scanner at 20X optical magnification. The scanned images were analysed as .VSI format files. Positive cell quantification in stroma and tumour cells were subsequently carried out using open source QuPath software [35] in a 10X field with standard settings for stains and with a threshold sensitivity of 0.3. The identification of tumour/stroma was done manually by a consultant histopathologist prior to quantification. Three separate representative fields per section were assessed for both tumour and stroma at 10X magnification.

### Isolation of stromal cells from primary multiple myeloma patient samples

Samples were collected and isolated by the Blood Cancer Network of Ireland, following informed consent and under an ethically approved protocol. Whole blood from primary multiple myeloma bone aspirates were centrifuged and supernatant was discarded. Red blood cells are lysed, and cells are plated in a T25 in α-MEM (Sigma, M4526) supplemented with 5% human serum, non-essential amino acids, sodium pyruvate, glutamax and penicillin-streptomycin. Stromal cell colonies form 4-7 days after isolation. Cells are trypsinised for 3 mins at 37°C, collected and plated for expansion. Cells are passaged twice before collection and analysis by flow cytometry.

### Human peripheral blood mononuclear cell (PBMC) isolation

PBMCs were isolated by density-gradient centrifugation from whole blood samples after written informed consent was obtained from healthy volunteers. Freshly drawn peripheral blood was collected in 5ml ethylene diamine tetra-acetic acid (EDTA) Vacutainer® tubes (BD Medical Supplies, Crawley, UK). PBMCs were isolated by layering 3ml of anti-coagulated blood over 3ml endotoxin-free Ficoll-Paque (Sigma-Aldrich) density-gradient solution in a 15ml tube (Sarstedt). Tubes were then centrifuged at 400 x g for 22 mins at 18°C with full acceleration (9) and brake off (0). Using a plastic Pasteur pipette (Sarstedt), the visible “buffy coat” layer of mononuclear cells was removed. PBMCs were transferred into fresh 15ml tubes (Sarstedt), washed twice with 10ml DPBS (ThermoFisher Scientific) and centrifuged at 400 x g for 5 mins at room temperature. The total number of live cells was determined by Trypan Blue (Sigma-Aldrich). Cells were resuspended in T cell media and added to co-culture assays..

### Human co-culture assays

For proliferation assays, PBMCs were isolated from healthy donor whole blood samples and single cell suspensions prepared as described above. Cells were re-suspended in complete medium (RPMI 1640 (ThermoFisher Scientific) supplemented with 10% heat-inactivated FBS, 1% sodium pyruvate, 1% non-essential amino acids, 1% L-glutamine, 1% penicillin/streptomycin and 0.1% β-mercaptoethanol (all Sigma-Aldrich)). Cells were stained with the CellTrace™ Violet proliferation kit (ThermoFisher Scientific) according to the manufacturer’s protocol and seeded in 96 well round bottom plates (Sarstedt) at a concentration of 1×10^5^ cells/100μl of complete medium with or without Human T-Activator CD3/CD28 Dynabeads® (ThermoFisher Scientific). NAFs/CAFs ± SI were added to wells of lymphocytes at a concentration of 1×10^4^ cells/100μl (ratio of 1:10 NAFs/CAFs:lymphocytes) of human complete MSC medium (RPMI 1640 medium (ThermoFisher Scientific) supplemented with 10% heat-inactivated FBS, 1% sodium pyruvate, 1% HEPES solution, 1% L-glutamine, 1% penicillin/streptomycin, 0.1% β-mercaptoethanol and 1ng/ml FGF2 (all Sigma-Aldrich)). After 96 hours, supernatants were and stored at -80°C. Cells were incubated with anti-human antibodies (see Table S1 for additional details) diluted in FACS buffer (DPBS supplemented with 1% FBS and 0.05% sodium azide (Sigma-Aldrich). Samples were analysed using a BD FACSCanto II Flow Cytometer (BD Biosciences). Flow cytometry data was analysed using FlowJo analysis software version 10 (Tree Star Inc.).

### Transcriptional data sets

Gene-expression profiles from two independent colorectal cancer data sets were accessed through the NCBI Gene-Expression Omnibus (http://www.ncbi.nlm.nih.gov/geo/) under accession numbers GSE35602 and GSE70468. GSE35602 contains microarray profiles separately profiled from laser-capture micro dissected stroma or epithelium regions from 13 colorectal cancer primary tumors (20). GSE70468 contains microarray profiles from primary fibroblasts derived using colon primary tumor and morphologically normal colonic mucosa tissue isolated from fresh colorectal cancer resection material. Both studies indicate that they were performed after approval by an institutional review board (IRB), and informed written consent was obtained from the subjects [36, 37].

### Bioinformatics analysis

Independent datasets were analysed using Partek Genomics Suite software (version 6.6; Partek Inc.). For the purpose of clustering, data matrices were standardized to the median value of probe set expression. Standardization of the data allows for comparison of expression for different probe sets. Following standardization, two-dimensional hierarchical clustering was performed (samples x probe sets/genes). Euclidean distance was used to calculate the distance matrix, a multidimensional matrix representing the distance from each data point (probe set-sample pair) to all the other data points. Ward’s linkage method was applied to join samples and genes together, with the minimum variance, to find compact clusters based on the calculated distance matrix.

### cBioPortal analysis

The cBioPortal for cancer genomics (http://www.cbioportal.org/) is an online open-access website for exploring, visualizing, and analyzing multidimensional cancer genomics data in the TCGA database [38, 39]. A total of 594 colorectal adenocarcinoma samples (TCGA, Pan Cancer Atlas) were analysed. The expression profiles of nine Siglec receptors and fibroblast activation protein (FAP) were investigated based on mRNA expression z-scores (RNA Seq V2 RSEM). Alterations in *siglec7* and *siglec9* and their association with overall survival (OS) is displayed as a Kaplan-Meier curve. The altered group contains patients that had *siglec7* and/or *siglec9* expression levels greater than 1.5 standard deviations above the average.

### Statistical analysis

Statistical analysis was performed using GraphPad® Version 9 (La Jolla, CA, USA). Experiments were performed in triplicate unless otherwise stated in figure legends, with two-three technical replicates per experiment. Data was assessed for normal distribution using D’Agostino-Pearson omnibus normality test. Datasets containing two groups were analyzed by unpaired t test, Mann-Whitney test or ratio paired t test where appropriate and indicated in the relevant figure legend. Datasets containing three or more groups were analyzed by ordinary one-way ANOVA followed by Tukey’s multiple comparisons test or two-way ANOVA with Sidak’s multiple comparisons test where appropriate and indicated. A p value of <0.05 was considered significant.

## Results

### Pro-inflammatory activation of stromal cells differentially regulates sialyltransferase expression resulting in increased cell surface α2,3- and α2,6-linked sialic acid

Mesenchymal stromal cells can sense and switch immune responses through secretion of soluble immunosuppressive molecules, as well as cell-cell contact mediated immunomodulatory ligand expression [15, 40]. MSC surface glycan expression affects their differentiation and homing ability; however, the impact of MSC glycosylation on their immunomodulatory potential remains unexplored. We therefore sought to assess the role of glycosylation on MSC-mediated suppression of T cell proliferation and activation. Using fluorescently labelled lectins, we assessed the levels of Con A, GNA, WGA and SNA-1 glycans by flow cytometry. A schematic depicting the preferential binding sites of each lectin is shown in Figure 1A. SNA-1 and WGA lectins, that bind sialic acid showed the highest level of expression on MSC when compared to control unstained MSC (Fig 1B). The two most common glycosidic linkages of sialic acid are α2,3 and α2,6 and therefore we next investigated the linkages of sialic acid on MSC and inflammatory activated MSC using the α2,3- and α2,6-binding lectins MAL-II and SNA-I, respectively. Inflammatory MSC expressed significantly higher levels of α2,3- and α2,6-linked sialic acids compared to control MSC (Fig 1C). To assess if alterations in sialic acid biosynthesis genes may be responsible for this increased sialylation, we analysed select sialic acid biosynthesis (Fig 1D), sialyltransferase (enzymes responsible for regulating sialylation) and sialidase (enzymes that cleave sialic acid) gene expression using RNA sequencing. Inflammatory MSC expressed similar levels of the sialylation biosynthesis enzymes GNE, NANP and CMAS, as well as the CMP-sialic acid transporter protein Slc35a1 (Fig 1C). We observed a significant reduction in NANS in inflammatory MSC but as the level of surface sialic acid was increased in these cells, the significance of this result is unclear (Fig 1C).

**Figure 1.**
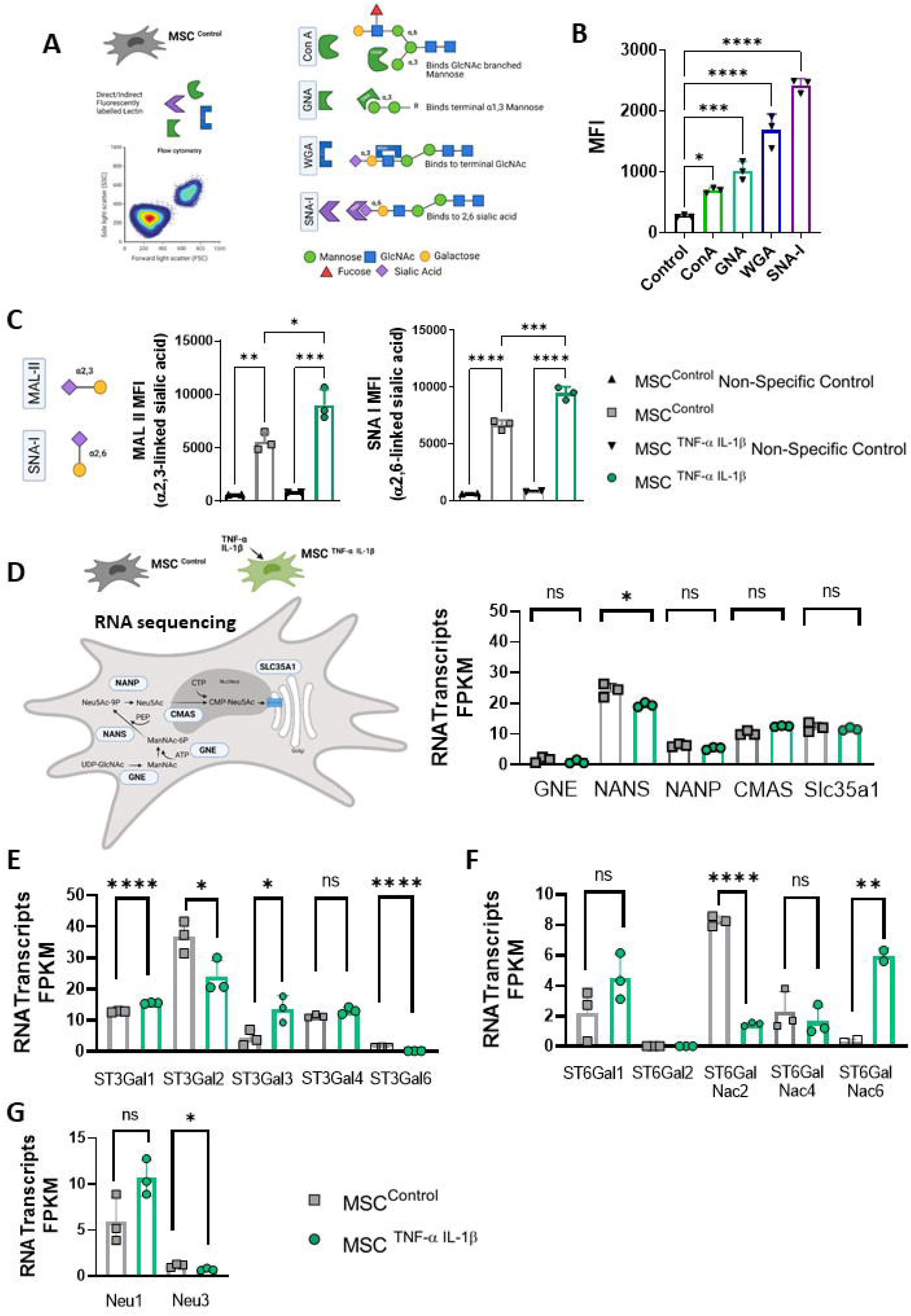
Pro-inflammatory cytokine activation induces sialylation on MSC. **(A)** Lectins with different sugar-binding preferences were screened for their ability to bind to Balb/c-derived MSC. **(B)** Median fluorescence intensity (MFI) of different biotinylated lectins bound to the MSC surface. Fluorescence was detected using an APC-conjugated streptavidin secondary antibody. **(C)** MFI of MAL II (α2,3-linked sialic acid) and SNA I (α2,6-linked sialic acid) expression on MSC^Control^ and MSC^TNF-α+IL-1β^. **(D)** Schematic depicting the key steps and genes involved in sialic acid biosynthesis and quantification of relevant RNA transcripts (Fragments Per Kilobase Million: FPKM) in MSC^Control^ and MSC^TNF-α+IL-1β^. **(E)** Quantification of α2,3-specific sialyltransferase RNA transcripts (FPKM) in MSC^Control^ and MSC^TNF-α+IL-1β^. **(F)** Quantification of α2,6-specific sialyltransferase RNA transcripts (FPKM) in MSC^Control^ and MSC^TNF-α+IL-1β^. **(G)** Quantification of neuraminidase (Neu1 and Neu3) RNA transcripts (FPKM) in MSC^Control^ and MSC^TNF-α+IL-1β^. Data are mean□±□SD; \**p*□<□0.05, \*\**p*□<□0.01, \*\*\**p*□<□0.001 and \*\*\*\**p* < 0.0001 using (**B** and **C**) one-way ANOVA with a Tukey post hoc test and (**D, E, F** and **G**) unpaired *t*-test. *n* = 2-3 biological replicates.

We next assessed mRNA expression levels of α2,3-linked specific (Fig 1E) and α2,6-linked specific (Figure 1F) sialyltransferases in inflammatory MSC. ST3Gal1 and ST3Gal3 are significantly increased in inflammatory MSC (Fig 1E). This would suggest that these two enzymes may be responsible for the increased α2,3-linked sialic acid linkages observed. Both ST3Gal1 and ST3Gal3 facilitate cancer cell immune evasion, adhesion, and metastasis in models of breast, ovarian and pancreatic cancer [41-43]. Similarly, for α2,6-specific sialyltransferases, our data showed that ST6Gal1 and ST6GalNac6 were more highly expressed in inflammatory MSC, suggesting a role in the higher levels of α2,6-linked sialic acid seen in Fig 1C. Sialidases, or neuraminidases, cleave terminal sialic acid residues from glycoproteins and glycolipids and contain four family members, Neu1-4 [44]. We observed no difference in expression of Neu1, however, expression of Neu3 were significantly reduced in inflammatory MSC (Fig 1G). This reduction in sialic acid-cleaving Neu3 may also contribute to the elevated levels of both α2,3- and α2,6-linked sialic acid observed on inflammatory MSC (Fig 1C). Both an increase in sialyltransferases and a reduction in sialidases in inflammatory MSC is associated with increased cell surface α-2,3 and α2,6 linked sialic acid expression.

### Pro-Inflammatory activated mesenchymal stromal cells suppress lymphocyte proliferation through a sialic acid-dependent mechanism

We have shown previously that inflammatory activated MSCs have potent immunosuppressive functions [45]. To test the impact of sialylation on inflammatory MSC immunosuppressive function, we pre-treated inflammatory MSCs with the sialyltransferase inhibitor (SI) 3F_ax_Neu5Ac, a sialic acid analogue [30]. Following treatment of pro-inflammatory MSC, we assessed MSC viability and phenotype (Fig 2A). SI treatment had no effect on MSC granularity or viability (Fig 2B), nor did it effect expression levels of the MSC characterisation markers CD44, CD73 and CD105 (Fig S1). Next, we assessed the effects of SI inhibition on cell surface α2,3 and α2,6 linked sialylation. We confirmed significant knockdown of MAL-II and SNA-I on inflammatory MSC following SI treatment (Fig 2C). These findings confirm that targeting sialyltransferase activity regulates the cell surface sialic acid expression in inflammatory MSC.

**Figure 2.**
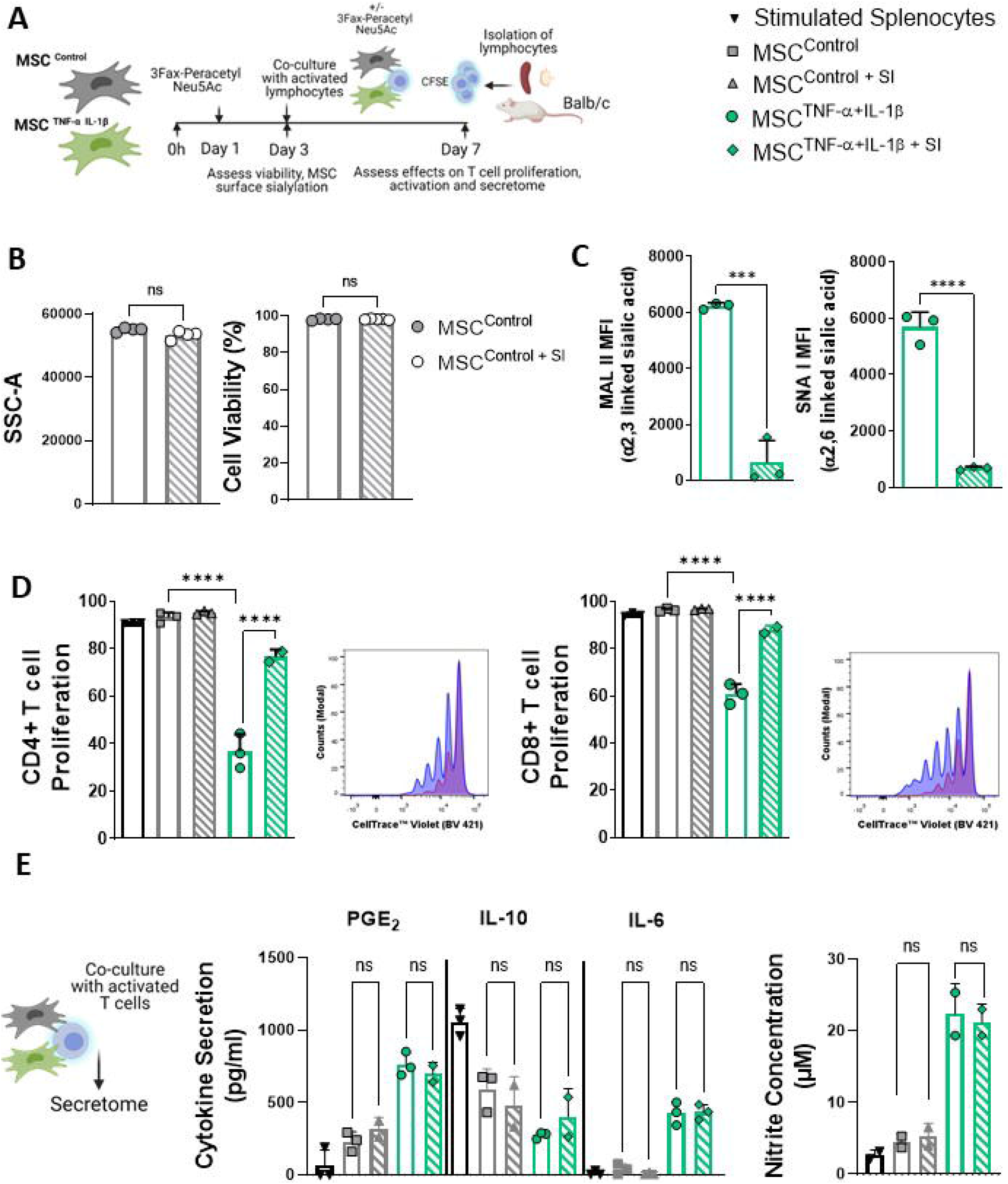
Enhanced suppressive ability of pro-inflammatory cytokine conditioned MSCs is dependent on sialylation. **(A)** Schematic overview of MSC-conditioning regime (± sialyltransferase inhibition (SI) using the sialic acid mimetic 3F_ax_Neu5Ac) and T cell co-culture assay set up. **(B)** Granularity and viability of MSC^Control^ with or without SI treatment measured by flow cytometry. **(C)** MFI of MAL II and SNA I binding on MSC^TNF-α+IL-1β^ with or without SI treatment measured by flow cytometry. **(D)** CD4+ and CD8+ T cell (CTV-labelled) proliferation following co-culture with MSC^Control^, MSC^TNF-α+IL-1β^ or MSC^TNF-α+IL-1β + SI^. Representative histograms depicting proliferation profiles of CD4+ and CD8+ T cells after co-culture with MSC^TNF-α+IL-1β^ or MSC^TNF-α+IL-1β + SI^ are also shown. **(E)** Following 96h co-culture, supernatants were collected and assayed for secreted levels of PGE_2_, IL-10 and IL-6 (ELISA) and for concentration of nitrites (Griess assay). Data are mean□±□SD; \*\*\**p*□<□0.001 and \*\*\*\**p* < 0.0001 using (**B, C** and **E**) unpaired *t*-test and (**D**) one-way ANOVA with a Tukey post hoc test. *n* = 2-4 biological replicates.

Using a syngeneic co-culture method (Fig 2A), we assessed the ability of inflammatory MSC to suppress CD4+ and CD8+ T cell proliferation. Following 96h co-culture, analysis of cell trace violet (CTV) profiles of both activated CD4+ and CD8+ T cells showed enhanced immunosuppression in co-cultures with TNF-α and IL-1β pre-activated MSC compared to untreated MSC controls (Figure 2D). Strikingly, SI treatment of inflammatory pre-activated MSC led to significant restoration of both CD4+ and CD8+ T cell proliferation (Fig 2D). Finally, we analysed co-culture supernatants for cytokines known to modulate pro-inflammatory activated MSC immunosuppression, including prostaglandin E2 (PGE2), IL-10, IL-6 and nitric oxide (NO). Pro-inflammatory activated MSC secreted significantly higher levels of soluble PGE2, NO and IL-6 (Fig 2E). However, we did not observe any significant differences in the levels of these soluble mediators in the presence or absence of the sialyltransferase inhibitor (Fig 2E) despite the restoration of T cell proliferation in these co-cultures. These findings show for the first time that inflammatory activated MSC-mediated suppression of T cell proliferation is dependent on MSC cell surface sialic acid-dependent interactions. This sialylation dependent immunosuppression may be independent of the secreted immunosuppressive factors PGE2 and NO.

### Inflammatory tumour secretome enhances the expression of α2,3 and α2,6-sialyl-transferases and α2,3 and/or α2,6 linked sialic acids in stromal cells which is associated with enhanced immunosuppression

Numerous studies have highlighted the role of stromal cells, including MSC and CAFs, in regulating immunity in solid and haematological tumours [15, 17]. Inflammatory tumour microenvironments can facilitate immunosuppression that enables tumour growth and progression [1]. CRC and MM arise in stromal dense microenvironments, where inflammation and stromal cell signatures are associated with immunosuppression [46]. MSC conditioned with tumour cell secretome (TCS) in the presence or absence of inflammation (iTCS) from multiple CRC and MM cell lines and models were assessed for sialyltransferase and sialic acid expression (Fig 3A). We observed increased levels of α2,6 linked SA on MSC^TCS^ and elevated levels of α2,3 linked SA observed on MSC^iTCS^ (Fig 3B). Furthermore, initial functional assessment revealed MSC^iTCS^ were significantly more suppressive than both MSC^Control^ and MSC^TCS^ at inhibiting CD8+ T cell proliferation (Figure 3B). Next, we utilised RNA sequencing analysis to identify potential changes in sialic acid biosynthesis and sialyltransferase expression at the transcriptional level. In contrast to inflammatory MSC, NANS expression was significantly increased in MSC^iTCS^ exclusively (Fig 3C). All other sialic acid biosynthesis genes screened were either unchanged or decreased in MSC^iTCS^. We show significantly increased expression of ST3Gal1 and ST3Gal3 (Fig 3D), α2,3-linkage specific sialyltransferase expression in MSC^iTCS^. These findings suggest that ST3Gal1 and ST3Gal3 may be crucial for stromal cell hyper-sialylation in the TME.

**Figure 3.**
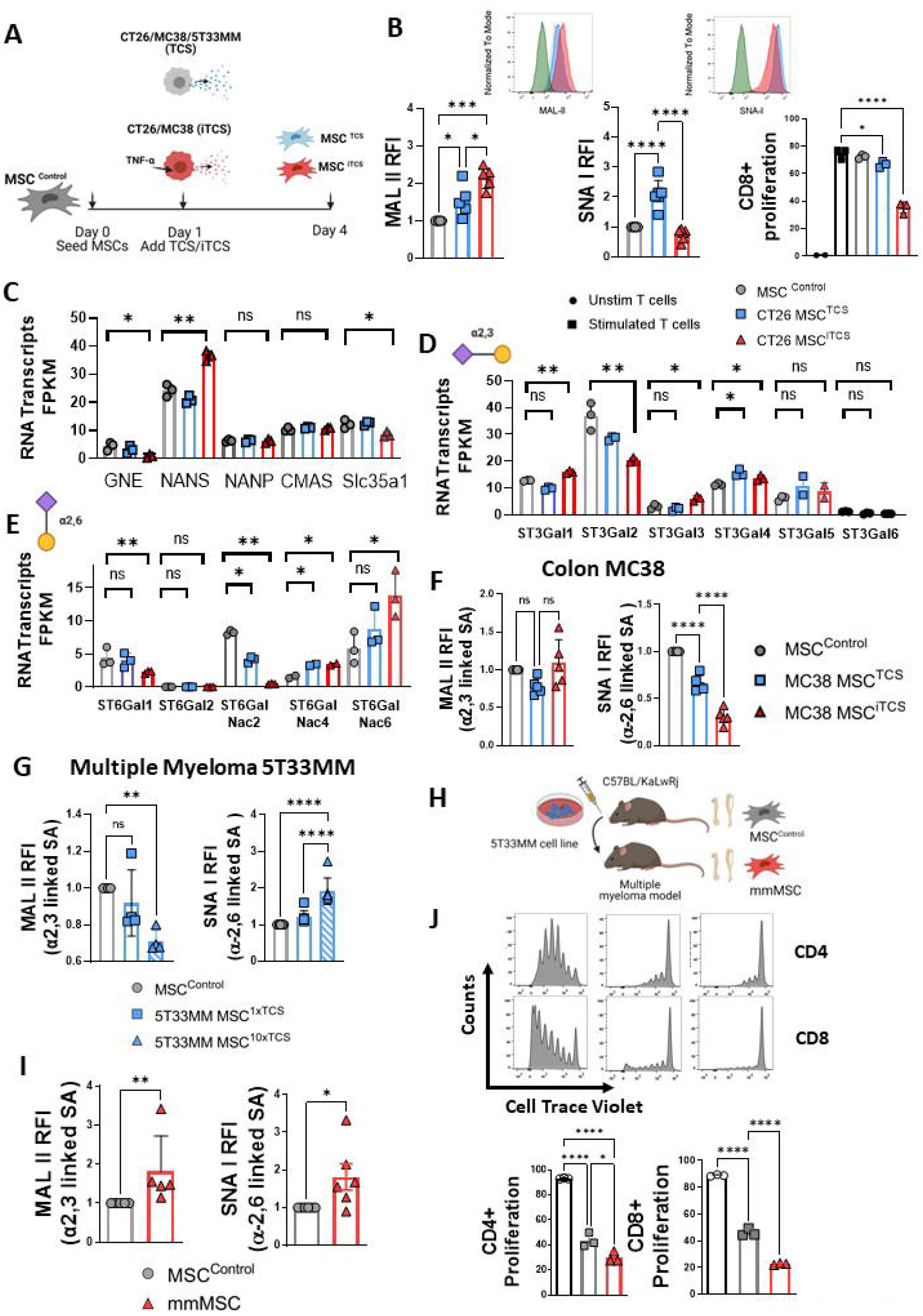
Inflammatory Tumor Conditioning leads to differential sialic acid expression on mouse MSCs. **(A)** Schematic overview of MSC-conditioning regime using tumour cell secretome (TCS) or inflammatory TCS (iTCS) from different mouse cancer cell lines (CT26 and MC38 = CRC; 5T33 = MM). **(B)** Relative fluorescence intensity (RFI: relative to MSC^Control^) of MAL II and SNA I binding on CT26 MSC^TCS^ and CT26 MSC^iTCS^ and representative histograms (inset). CD8+ T cell proliferation following co-culture with MSC^Control^, CT26 MSC^TCS^ or CT26 MSC^iTCS^. **(C)** Quantification of key sialic acid biosynthesis-related RNA transcripts (FPKM) in MSC^Control^, CT26 MSC^TCS^ and CT26 MSC^iTCS^. **(D)** Quantification of α2,3-specific sialyltransferase RNA transcripts (FPKM) in MSC^Control^, CT26 MSC^TCS^ and CT26 MSC^iTCS^. **(E)** Quantification of α2,6-specific sialyltransferase RNA transcripts (FPKM) in MSC^Control^, CT26 MSC^TCS^ and CT26 MSC^iTCS^. **(F)** RFI (relative to MSC^Control^) of MAL II and SNA I binding on MC38 MSC^TCS^ and MC38 MSC^iTCS^. **(G)** RFI (relative to MSC^Control^) of MAL II and SNA I binding on 5T33MM MSC^1xTCS^ and 5T33MM MSC^10xTCS^. **(H)** Schematic overview illustrating induction of 5T33 murine MM model. **(I)** RFI (relative to MSC^Control^) of MAL II and SNA I binding on MM-derived MSC. **(J)** CD4+ and CD8+ T cell proliferation following co-culture with MSC^Control^ or MM-derived MSC. Representative histograms depicting proliferation profiles of CD4+ and CD8+ T cells after co-culture with MSC^Control^ or MM-derived MSC are also shown. Data are mean□±□SD; \**p*□<□0.05, \*\**p*□<□0.01, \*\*\**p*□<□0.001 and \*\*\*\**p* < 0.0001 using (**B, F, G** and **J**) one-way ANOVA with a Tukey post hoc test, (**C, D** and **E**) unpaired *t*-test and (**I**) Mann-Whitney test. *n* = 2-6 biological replicates.

The expression pattern of α2,6 sialyl-transferases were also differentially regulated in the inflammatory tumour microenvironment (Fig 3E). ST6GalNac 4 and 6 were significantly upregulated on MSC^TCS^ and MSC^iTCS^ when compared to MSC^Control^ (Fig 3E). ST6GalNac1, 2 wer downregulated in MSC^iTCS^ compared to MSC^Control^ or MSC^TCS^, ST6GalNac1, 3 and 5 were not detected in our RNA seq dataset. This may suggest that the increased α2,6 sialic acid on MSC^TCS^ is regulated by a balance of sialyltransferase activity and sialidase activity in the TME. In the MC38 CRC model, α2,3 sialic acid expression was increased on MSCs^iTCS^ (p=.056) compared to control MSC or MSC^TCS^. However, MC38 MSC^TCS^ or MSC^iTCS^ expressed lower levels of α2,6 sialic acid than MSC^Control^ (Fig 3F) These results suggest although the overall level of sialylation is high, stromal cell sialic acid expression may be tumour microenvironment-specific and differentially regulation may dependent on discrete tumour lines genetic features and characteristics. Similar observations were noted in the 5T3MM model, of MM. We conditioned MSC with the secretome from ex vivo 5T33MM multiple myeloma (MM) murine cells at two concentrations of 5T33MM TCS, 1x and 10x. We observed a dose-dependent decrease in α2,3 and α2,6 sialic acid expression on MSC^TCS^ (Fig 3G). To assess if ex vivo conditioning recapitulated the in vivo TME, we isolated and expanded stromal cells directly from the bone marrow of diseased 5T33MM mice (mmMSC) and assessed their sialic acid expression compared to wild type C57BL/KaLwRj-derived MSC (MSC^Control^) (Fig 3H). Both α2,3 (MAL-II) and α2,6 (SNA-I) sialic acid expression was significantly higher on MSC derived from 5T33MM mice (Fig 3I). The 5T33MM-derived MSC were also more suppressive than their WT-derived controls and could significantly inhibit both allogeneic CD4+ and CD8+ T cell proliferation compared to control (Fig 3J). Taken together, these data confirm that stromal cells in the two inflammatory tumour microenvironments express higher baseline levels of α2,3 and/or α2,6 sialic acid and is associated with enhanced immunosuppression.

### Inflammatory tumour secretome enhances stromal cell-mediated suppression of T cell proliferation and activation, which is reversed by inhibiting sialyl-transferase activity

Although increases in sialo glycan density or hyper sialylation in tumour cells has been observed, we show here for the first time that stromal cell sialylation can be modulated by the inflammatory tumour microenvironment. To determine the significance of sialylation of stromal cells in the TME, we assessed sialic acid expression of tumour conditioned stromal cells compared to colon cancer epithelial cells. Strikingly, while MSC^iTCS^ and MSC^TCS^ expressed the highest of α2,3 and α2,6 sialic acid, respectively, baseline levels of sialic acid were higher in stromal cells compared to CT26 epithelial cells (Fig 4B). To assess the binding affinities of these sialylated proteins for Siglec receptors, we used a Siglec E receptor Fc chimera [47]. Siglec E, a homologue of human Siglec 7/9, is expressed by immune cells such as T cells, macrophages and neutrophils [27]. It contains an inhibitory ITIM (immunoreceptor tyrosine-based inhibitory motif) in its cytoplasmic domain [48]. Siglec E ligand on CT26 and MC38 conditioned stromal cells was differentially induced by the tumour cell secretome, in the presence or absence of inflammation (Fig 4C, left, middle). Similar to sialic acid expression shown in Fig 4B, both control and tumour conditioned stromal cells had significantly higher levels of Siglec E ligand than cancer cells (Fig 4C, right). Our results show that sialic acid expression and Siglec E ligand expression are higher on stromal cells than tumour cells in the TME.

**Figure 4.**
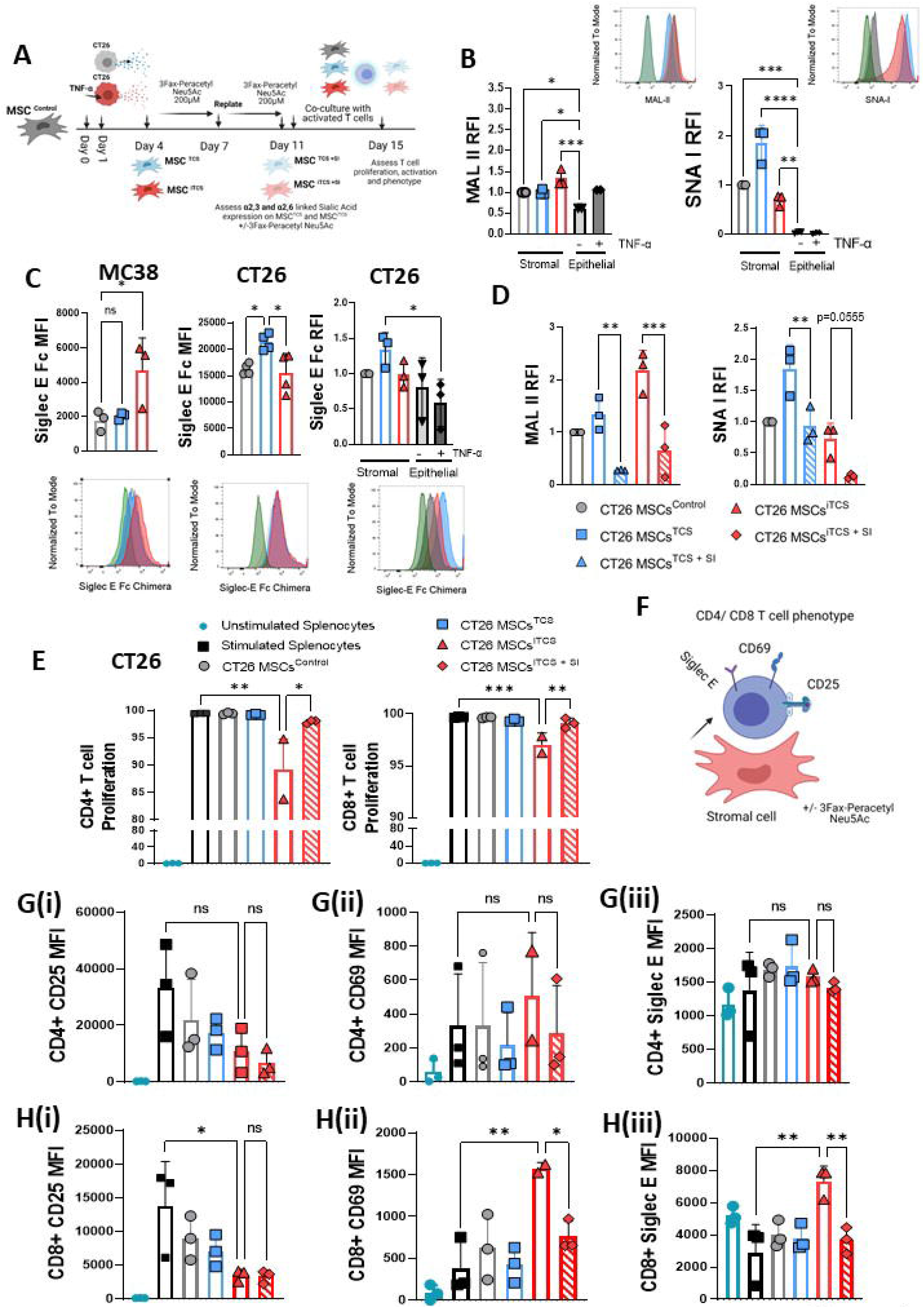
Inflammatory tumour-conditioned MSCs suppress T cell effector phenotype in a sialylation-dependent manner. **(A)** Schematic overview of MSC-conditioning regime (± SI pre-treatment) using TCS or iTCS and T cell co-culture assay set up. **(B)** RFI (relative to MSC^Control^) of MAL II and SNA I expression on CT26 MSC^TCS^, CT26 MSC^iTCS^ and CT26 cancer cells ± TNF-α conditioning, and representative histograms (inset). **(C)** MFI or RFI (relative to MSC^Control^) of Siglec E Fc chimera expression on MSC^Control^, MC38 MSC^TCS^ and MC38 MSC^iTCS^ or MSC^Control^, CT26 MSC^TCS^, CT26 MSC^iTCS^ and CT26 cancer cells ± TNF-α conditioning, and representative histograms. **(D)** RFI (relative to MSC^Control^) of MAL II and SNA I expression on CT26 MSC^TCS^, CT26 MSC^TCS + SI^, CT26 MSC^iTCS^ and CT26 MSC^iTCS + SI^. **(E)** CD4+ and CD8+ T cell proliferation following co-culture with MSC^Control^, CT26 MSC^TCS^, CT26 MSC^iTCS^ or CT26 MSC^iTCS + SI^. **(F)** Schematic depicting T cell expression of activation (CD25), inhibitory (CD69) and immunomodulatory (Siglec E) markers following exposure to conditioned MSCs. MFI of CD25 **(Gi)**, CD69 **(Gii)** and Siglec E **(Giii)** on CD4+ T cells following co-culture with MSC^Control^, CT26 MSC^TCS^, CT26 MSC^iTCS^ or CT26 MSC^iTCS + SI^. MFI of CD25 **(Hi)**, CD69 **(Hii)** and Siglec E **(Hiii)** on CD8+ T cells following co-culture with MSC^Control^, CT26 MSC^TCS^, CT26 MSC^iTCS^ or CT26 MSC^iTCS + SI^. Data are mean□±□SD; \**p*□<□0.05, \*\**p*□<□0.01, \*\*\**p*□<□0.001 and \*\*\*\**p* < 0.0001 using (**B, C, D, E, G** and **H**) one-way ANOVA with a Tukey post hoc test. *n* = 2-4 biological replicates.

We next assessed the functional effects of de-sialylation of tumour conditioned stromal cells on T cell function and phenotype. The optimum concentration of sialyl-transferase inhibitor 3Fax5NeuAc, (SI) was determined in vitro (Fig S2). SI pre-treatment revealed no significant differences in cell viability, granularity, size or morphology (Fig S3A-D). SI treatment led to significant reductions in both SNA-I and MAL-II expression on TCS or iTCS treated MSC (Fig 4D). Tumour conditioned stromal cells were pre-treated with SI and subsequently co-cultured with syngeneic lymphocytes (Fig 4A). As shown in Fig 4E, iTCS-conditioned MSC significantly inhibited both CD4+ and CD8+ T cell proliferation. SI treatment abolished this effect with significant restoration of proliferation observed for both T cell subsets. These results show that sialylation plays a key role in tumour induced MSC-mediated immunosuppression. We next assessed T cell activation and phenotype, including CD25, CD69 and Siglec E expression (Fig 4F). We observed that MSC^iTCS^ significantly inhibited CD25 expression on CD8+ T cells (Fig 4Hi), with a clear trend towards suppression on CD4+ T cells (Fig 4Gi). CD25 expression was unaffected by inhibition of stromal cell sialylation (Fig 4Gi + Hi). CD8+ T cells co-cultured specifically with MSC^iTCS^ expressed significantly higher levels of CD69 (Fig 4Hii). Targeting sialyltransferase activity with SI in MCS^iTCS^ significantly reduced the level of CD69 expression on CD8+ T cells but not on CD4+ T cells (Fig 4Hii and 4Gii). MSC^iTCS^ induced elevated levels of Siglec E expression on CD8+ T cells specifically (Fig 4Hiii). This increased expression was significantly abrogated following inhibition of stromal cell sialyl-transferase activity (Fig 4Hiii). In contrast, Siglec E expression on CD4+ T cells was unaffected by incubation with iTCS-conditioned MSC in the presence or absence of SI (Fig 4Giii). CD8+ T cell phenotype was most significantly altered following inhibition of sialyltransferase activity in tumour conditioned stromal cells. We have shown here for the first time that stromal cell sialylation is induced following tumour conditioning which enhances stromal cell immunosuppression and can dictate CD8+ T cell phenotype, Siglec expression and function.

### Human tumour conditioned MSC and stromal cells in CRC tumours express higher levels of ST enzyme, higher levels of sialic acid and siglec ligand expression than epithelial cells in the TME

To assess the clinical relevance of these findings, we assessed stromal cell and epithelial cell areas in human histopathological specimens. Assessment of T cell localisation with stromal or epithelial areas in CRC revealed a significantly higher density of CD3 and CD8 T cells with stromal regions in CRC than with epithelial cells (Fig 5A, B). Quantification using QuPath pathology software validated a significantly higher number of CD3 (Fig 5C, upper) and CD8 T cells (Fig 5C, lower) associated with stromal regions than epithelial regions in colorectal tumours. Next we used human data sets, human tumour cell lines and stromal cells to investigate the expression of multiple α2,3-and α2,6-specific sialyltransferases in human colorectal cancer tissue. We analysed gene expression profiles (GEO accession no. GSE35602) of colorectal cancer resection samples which had been laser-capture micro-dissected to separate the stromal and epithelial fractions prior to microarray profiling [36]. Assessment of multiple sialyltransferases within this dataset indicated that expression of the α2,3-specific sialyltransferases ST3Gal1, ST3Gal4 and ST3Gal6 more closely associated with the stromal compared to the epithelial compartment as evidenced by co-expression of the mesenchymal lineage markers α-SMA (ACTA2), vimentin (VIM), CD90 (THY1), PDGFR-α (PDGFRA) and CD105 (ENG) (Fig 5D) A) [15]. Quantification of relative gene expression of ST3Gal1, ST3Gal4 and ST3Gal6 confirmed this, showing significantly higher expression of all three genes in the stromal fraction compared to the epithelial fraction (Fig 5E). Expression the α2,6-specific sialyl-transferases ST6Gal1 and ST6Gal2 were not significantly altered (Fig 5E). Additionally, we observed significantly higher expression of α2,6-specific ST6GALNAC6, which preferentially adds sialic acid to glycolipids as opposed to glycoproteins, along with a significant decrease in the sialidase NEU1 (Fig 5F). To extend these observations, we analysed transcriptional profiles (GEO accession no. GSE70468) of patient-matched primary fibroblasts (n = 14 samples from 7 patients) isolated from within colorectal cancer tissue (cancer-associated fibroblasts – CAFs) and from adjacent, normal mucosal tissue (normal-associated fibroblasts – NAFs) [37]. While we observed no significant differences in expression levels of ST3Gal1, ST3Gal4 or ST3Gal6 between NAFs and CAFs, relative expression levels of both α2,6-linkage specific sialyltransferases (ST6Gal1 and ST6Gal2) were significantly increased in CAFs compared to patient-matched NAFs (Fig 5G). Taken together, this data suggests that the stromal compartment of the colorectal cancer microenvironment has elevated levels of α2,3-linkage specific sialyltransferases compared to the epithelial compartment but, additionally, within the stromal compartment, tumour-educated fibroblasts (CAFs) express higher levels of α2,6-linkage specific sialyltransferases compared to normal adjacent stromal cells.

**Figure 5.**
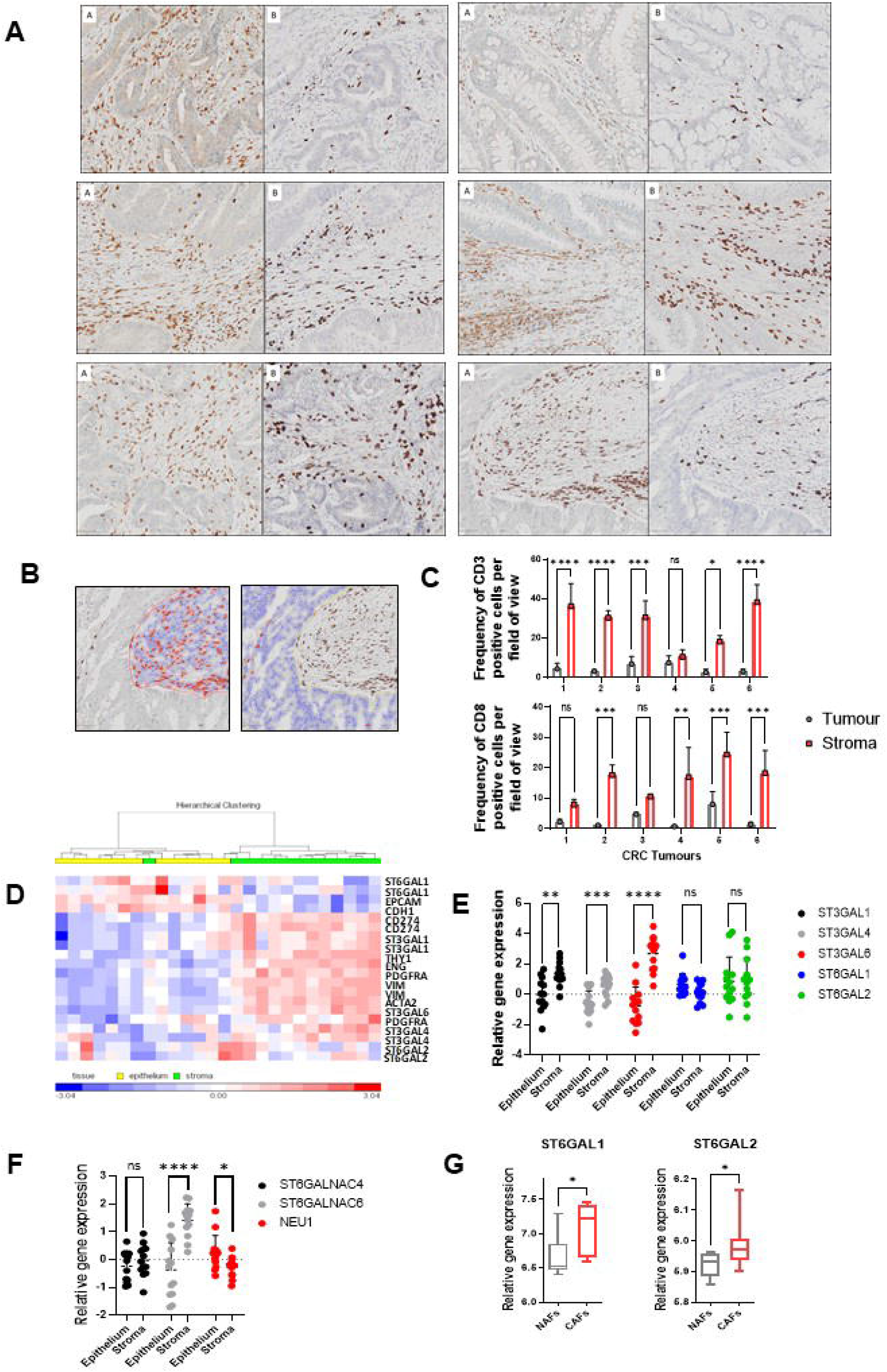
CRC tumour-associated stroma has higher T cell infiltration and levels of ST gene expression compared to tumour epithelium. (A) Representative immunohistochemical stained sections of 6 individual CRC patient tumours. Sections in A panels are stained with anti-CD3 (brown) while sections in B panels are stained with anti-CD8 (brown) (n=6). (B) Representative section demonstrating positive cell quantification analysis for CD3+ cells carried out using open source QuPath software for both stroma (left) and tumour (right). Positive cells are detected with a red outline and negative cells are blue. (C) Bar graphs showing frequency % of CD3+ cells (top panel) and CD8+ T cells (bottom panel) per field of view. (D) Clustering for gene-expression profiles of PDGFR-α (PDGFRA), PD-L1 (CD274), CD90 (THY1), CD105 (ENG), Vimentin (VIM), a-SMA (ACTA2), E-cadherin (CDH1), EpCAM (EPCAM), ST3Gal1, ST3Gal4, ST3Gal6, ST6Gal1 and ST6Gal2. Relative gene-expression of (E) ST3Gal1, ST3Gal4, ST3Gal6, ST6Gal1 and ST6Gal2 and (F) ST6GalNac4 and ST6GalNac6 by epithelial and stromal cells from colorectal cancer patients (n = 13; data set GSE35602). (G) Relative gene-expression of ST6Gal1 and ST6Gal2 on normal-associated fibroblasts (NAFs) and cancer-associated fibroblasts (CAFs) (n = 7; data set GSE70468). Data are mean□±□SD; *p□<□0.05, **p□<□0.01, ***p□<□0.001 and ****p < 0.0001 using (C) two-way ANOVA with Sidak’s multiple comparisons test and (E, F and G) Mann-Whitney test.

Using TCS or iTCS from two human colorectal cancer cell lines, HT29 and HCT116, we then conditioned human bone marrow derived MSC (Fig 6A) and analysed α2,3 and α2,6 sialic acid expression. Similar to observations in murine models, baseline sialylation levels were significantly higher on stromal cells compared to epithelial cells (Fig 6B & C). Expression of α2,3 sialic acid expression on stromal cells is differentially expressed following conditioning with HT29 and HCT116 TCS/iTCS (Fig 6B and C left panels). We observed significantly increased α2,6 sialic acid expression on HT29 MSC^iTCS^ and a trend in increase on HCT116 conditioned stromal cells (Fig 6B and C left panels). We further characterised the effects of TCS/iTCS-conditioning on MSC using a different tumour model, multiple myeloma. TCS/iTCS from two MM tumour lines RPMI 8226 and MM1S was generated and harvested as for CRC models. α2,3 and α2,6 sialic acid expression on stromal cells conditioned with RPMI 8226 TCS/iTCS or MM1S TCS/iTCS showed the same trend as observed for HT29-conditioned MSC, with α2,3 sialic acid expression comparable between groups and α2,6 sialic acid expression increased following stromal cell iTCS-conditioning (Fig S4A and B). Importantly, this same trend in expression of α2,3 and α2,6 sialic acid was observed in multiple myeloma patient derived MSC compared to healthy donor bone marrow (Fig S4C). Taken together, these findings provide clues on the potential Siglec receptors that may be important in regulating stromal-immune cell interactions within the TME based on their sialic acid binding preferences.

**Figure 6.**
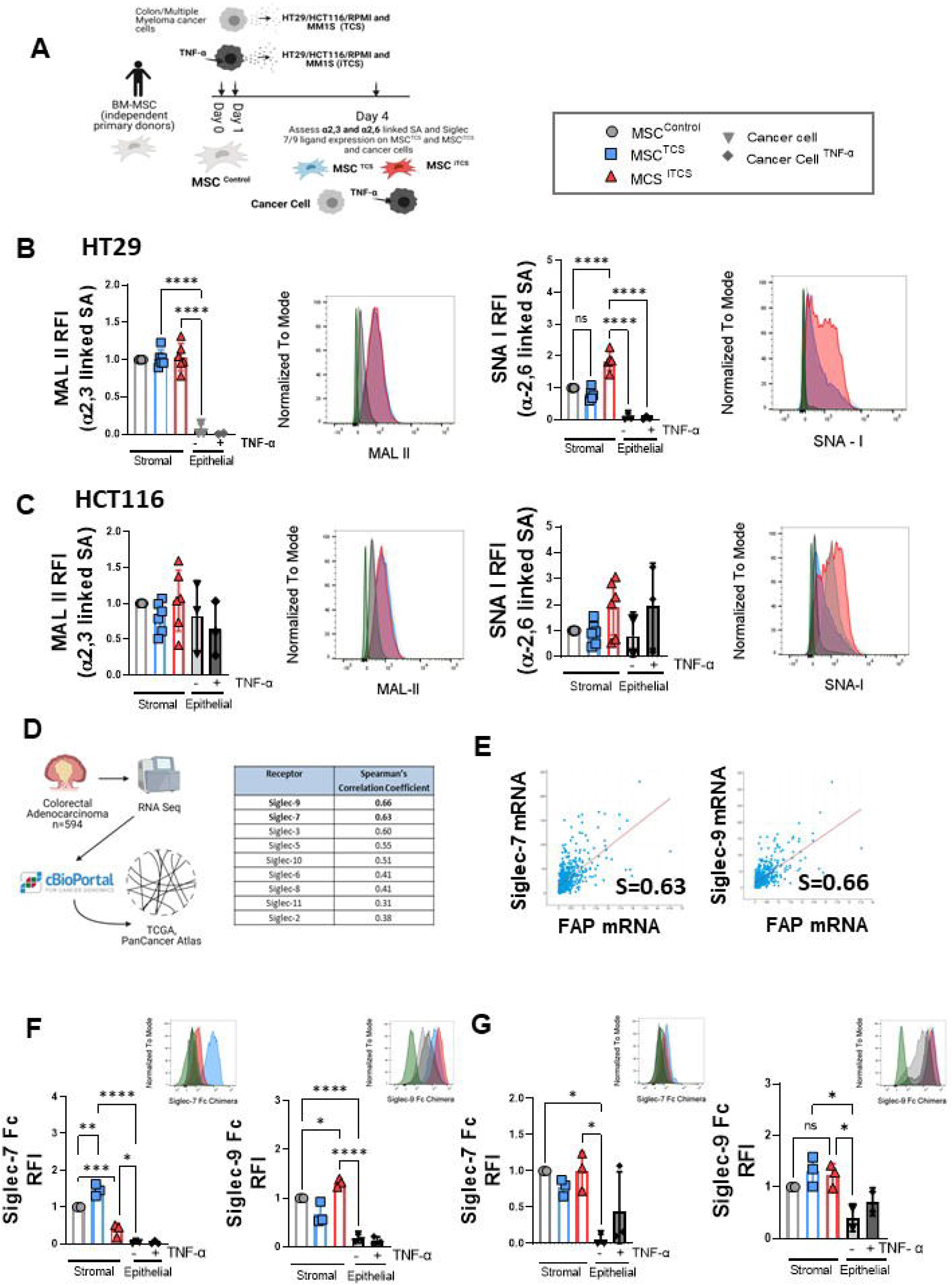
Tumor conditioned human MSCs have elevated levels of sialic acid expression. **(A)** Schematic overview of human bone marrow-derived MSC-conditioning regime using TCS or iTCS from two CRC cell lines, HT29 and HCT116. **(B)** RFI (relative to MSC^Control^) of MAL II and SNA I expression on HT29 MSC^TCS^, HT29 MSC^iTCS^ and HT29 cancer cells ± TNF-α conditioning, and representative histograms. **(C)** RFI (relative to MSC^Control^) of MAL II and SNA I expression on HCT116 MSC^TCS^, HCT116 MSC^iTCS^ and HCT116 cancer cells ± TNF-α conditioning, and representative histograms. **(D)** cBioPortal was used to search TCGA database for colorectal cancer patient cohorts and for analysis of ITIM-containing Siglec receptor relative gene expression. **(E)** Dot plots showing the correlation between FAP and Siglec-7 or -9 receptor expression in a cohort of 594 CRC patient samples (S = Spearman’s correlation coefficient). **(F)** RFI (relative to MSC^Control^) of Siglec-7 and Siglec-9 expression on HCT116 MSC^TCS^, HCT116 MSC^iTCS^ and HCT116 cancer cells ± TNF-α conditioning, and representative histograms (inset). **(G)** RFI (relative to MSC^Control^) of Siglec-7 and Siglec-9 expression on HT29 MSC^TCS^, HT29 MSC^iTCS^ and HT29 cancer cells ± TNF-α conditioning, and representative histograms (inset). Data are mean□±□SD; \**p*□<□0.05, \*\**p*□<□0.01, \*\*\**p*□<□0.001 and \*\*\*\**p* < 0.0001 using **(B, C, F** and **G)** one-way ANOVA with a Tukey post hoc test. *n* = 2-6 biological replicates.

To explore associations between stromal cells and Siglec receptors, we next accessed the cBioPortal tool to evaluate potential correlations between fibroblast activation protein (FAP), as a marker for stromal cells, and each of the nine ITIM motif carrying Siglec receptors known to be expressed by immune cells in humans (Fig 6D). A total of 594 patients in the Colorectal Adenocarcinoma TCGA PanCancer Atlas dataset were analysed. As shown in Fig 6D,E, the two Siglec receptors with the strongest positive correlation with FAP were Siglec 9 and 7 (0.66 and 0.63, respectively), as determined by Spearman’s correlation coefficient. Therefore, we assessed human bone marrow derived MSC ± HT29 and HCT116 TCS/iTCS conditioning, for expression of specific Siglec 7/9 ligands using Siglec 7/9 Fc chimeras. HCT116 TCS-conditioned MSC expressed the highest level of Siglec 7 ligand, while HCT116 iTCS-conditioned MSC expressed significantly higher levels of Siglec 9 ligand (Fig 6F). HT29 TCS/iTCS-conditioned stromal cells expressed comparable levels of Siglec 7 and 9 ligands (Fig 6G), and stromal cells with/without tumour cell conditioning expressed higher levels of siglec ligands than cancer cells (Fig 6F &G).

### CAFs have enhanced sialic acid/Siglec ligand expression and have a more potently immunosuppressive phenotype that is associated with induction of a Siglec receptor-expressing exhausted phenotype in CD8+ T cells

Next, we elucidated whether altered expression of Siglec 7 and/or 9 had prognostic implications in CRC. Analysis of a cohort of 522 patient samples from the Colorectal Adenocarcinoma TCGA PanCancer Atlas dataset of which 46 had altered expression of one or both of Siglec 7 and/or 9 >1.5 standard deviations (SDs) above the average revealed a clear trend (p=0.0594) towards worse overall survival (Fig 7A). We next sought to investigate stromal cell sialylation in clinical CRC specimens. CAFs were isolated from CRC tumours, and patient-matched cancer-associated normal fibroblasts (NAFs) were isolated and cultured from tumour-adjacent non-cancerous tissue (Fig 7B). NAFs and CAFs were analysed for expression of typical stromal cell characterisation markers (Fig S5). Using lectin-based flow cytometry, we observed that both NAFs and CAFs expressed α2,3 sialic acid at comparable levels, however, α2,6 sialic acid expression was significantly higher in CAFs (Fig 7C). Specific Siglec ligand staining revealed that, while Siglec 7 ligand was expressed by both NAFs and CAFs, Siglec 9 ligand was significantly higher on CAFs (Fig 7D). To assess the functional effects of sialylation on stromal cell-mediated immunosuppression, we co-cultured CRC patient-derived NAFs and CAFs with donor PBMCs. Both stromal cell populations could significantly suppress CD8+ compared to anti-CD3/CD28 stimulated PBMCs alone, CAFs were significantly more suppressive than NAFs (Fig 7E). Furthermore, CAFs could induce a significantly higher proportion of CD8+ T cells with a more exhausted phenotype than NAFs, as characterised by CD69 expression [49] (Fig 7F) and CD69/Tim-3 co-expression (Fig 7G). Frequencies of CD4+CD69+ and CD4+CD69+Tim-3+ T cells were not significantly increased after co-culture with CAFs (Fig S7B & C), indicating these CAF-mediated effects were CD8+ T cell-specific. While individual expression of an inhibitory receptor does not necessarily indicate exhaustion, co-expression of multiple inhibitory receptors is a principal feature of exhausted T cells [50].

**Figure 7.**
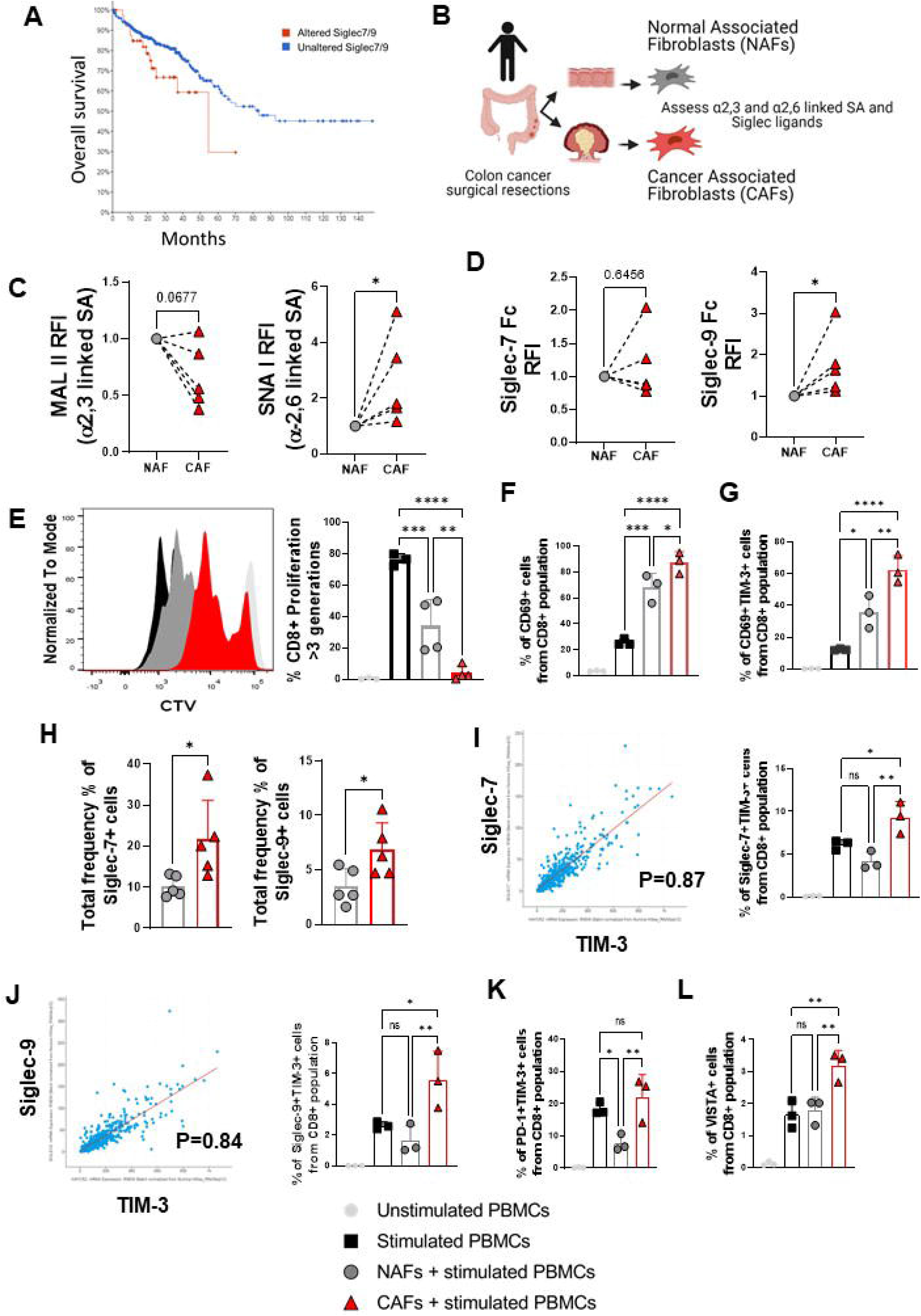
CRC tumour-derived CAFs have elevated sialic acid expression and induce an exhausted phenotype in CD8+ T cells. **(A)** cBioPortal analysis of the same CRC patient dataset used in Figure 6E showing overall survival (months) in patients with altered Siglec-7 and/or Siglec-9 gene expression vs those with unaltered expression. **(B)** Schematic overview depicting the tissue from which NAFs and CAFs were isolated. **(C)** RFI (relative to NAFs) of MAL II and SNA I expression on CAFs. **(D)** RFI (relative to NAFs) of Siglec-7 and Siglec-9 Fc chimera expression on CAFs. **(E)** CD8+ T cell proliferation following co-culture with NAFs or CAFs. Representative histograms depicting proliferation profiles of CD8+ T cells after co-culture with NAFs or CAFs are also shown. **(F)** Frequency (%) of CD8+CD69+ T cells after co-culture with NAFs or CAFs. **(G)** Frequency (%) of CD8+CD69+Tim-3+ T cells after co-culture with NAFs or CAFs. **(H)** Frequency (%) of all Siglec-7 and Siglec-9-expressing cells after co-culture with NAFs and CAFs. **(I)** Dot plots showing the correlation between Siglec-7 and Tim-3 expression in a cohort of 594 CRC patient samples (S = Spearman’s correlation coefficient) and frequency (%) of CD8+Siglec-7+Tim-3+ T cells after co-culture with NAFs or CAFs. **(J)** Dot plots showing the correlation between Siglec-9 and Tim-3 expression in a cohort of 594 CRC patient samples and frequency (%) of CD8+Siglec-9+Tim-3+ T cells after co-culture with NAFs or CAFs. Frequency (%) of **(K)** PD-1+Tim-3+ and **(L)** VISTA+ CD8+ T cells after co-culture with NAFs or CAFs. Data are mean□±□SD; \**p*□<□0.05, \*\**p*□<□0.01, \*\*\**p*□<□0.001 and \*\*\*\**p* < 0.0001 using **(C** and **D)** ratio paired t test and **(E, F, G, H, I, J, K** and **L)** one-way ANOVA with a Tukey post hoc test. *n* = 3-5 biological replicates.

As Siglec 7 and 9 are inhibitory receptors, we assessed cBioPortal data for correlations between both Siglecs and Tim-3 in the same Colorectal Adenocarcinoma TCGA PanCancer Atlas dataset as utilised in Figure 7I. The results showed that both Siglec 7 and 9 had a strong positive correlation with Tim-3 (0.87 and 0.84, respectively), as determined by Spearman’s correlation coefficient (Fig 7I and J). We show that following co-culture with CA, but not NAFs, the frequency of Siglec 7 and siglec 9 receptor expressing cells is significantly increased (Fig 7H, left and right). Importantly, we also observed significant increases in the frequencies of both Siglec 7+Tim-3+ and Siglec 9+Tim-3+ CD8+ T cells following co-culture with CAFs specifically (Fig 7I and J). To validate these CAF-mediated effects on CD8+ T cells, we analysed the co-cultured T cells for expression of the additional negative checkpoint regulators (NCRs) PD-1 and VISTA. The frequency of PD-1+Tim-3+ co-expressing CD8+ T cells was significantly increased following co-culture with CAFs (Fig 7K), as was the frequency of VISTA-expressing CD8+ T cells (Fig 7L). We observed no significant increase in frequencies of either PD-1+, PD-1+Tim-3+ co-expressing or VISTA+ CD4+ T cells following co-culture with CAFs (Fig S7D, E and F, respectively). This expansion of NCR-expressing CD8+ T cells was not evident following co-culture with patient-matched NAFs. These data provide strong evidence that CAFs induce an exhausted CD8+ T cell phenotype than NAFs and that inhibitory Siglecs such as Siglec 7 and 9 may be indicative of this phenotype and considered alongside more established T cell markers like PD-1, LAG-3, VISTA and CTLA4 when defining T cell exhaustion.

### CAF induction of exhausted CD8+ T cells is reversible by targeting sialyl-transferase enzyme activity using 3F_ax_Neu5Ac

We next assessed the effect(s) of de-sialylation on CAF-induced CD8+ T exhaustion. NAFs/CAFs were treated with the SI 3F_ax_Neu5Ac prior to co-culture with T-cells (Fig 8A). We confirmed significant inhibition of Siglec 9 ligand expression on SI-treated CAFs (Fig S6). Following co-culture, we analysed the frequencies of Siglec 7- and 9-expressing CD8+ T cells. As can be seen in Fig 8B (left) and C (left), levels of Siglec 7 and Siglec 9 receptor-expressing CD8+ T cells, respectively, were significantly increased after co-culture with CAFs specifically. Furthermore, CAFs also induced a significantly higher percentage of CD8+PD-1+ T cells (Fig 8D (left panel). Importantly, pre-treatment of CAFs with SI prior to addition to the co-culture resulted in significant decreases in the frequencies of both Siglec-7- and 9-expressing CD8+ T cells (Fig 8B, (right panel) and C (right panel)). This effect was specific to CAFs, as frequencies of CD8+ T cell populations were unchanged following co-culture with NAFs +/- SI pre-treatment (Fig 8B, (middle panel) and C (middle panel)). CAFs also induced higher frequencies of Siglec 9-expressing CD4+ T cells (Fig S7G(i)) but, in contrast to its effect on CD8+ T cells, SI pre-treatment had no effect on reversing this increase (Fig S7G(ii) and (iii)). We observed a clear trend towards reduction in the frequency of CD8+PD-1+ T cells (Fig 8D (right panel)) in co-cultures with SI treated CAFs. This effect was CAF specific as there was no significant effect in SI treated NAFs (Fig 8D (middle panel). These results demonstrate that TME-derived CAFs can suppress activated T-cells and promote CD8+ T cell exhaustion and that this immunosuppressive effect is significantly reversed through the modulation of sialylation on the stromal cell surface. Understanding how sialylation of stromal cells is regulated and functions to enhance immunosuppression in the TME could uncover novel immune checkpoints to reactivate anti-tumour immunity.

**Figure 8.**
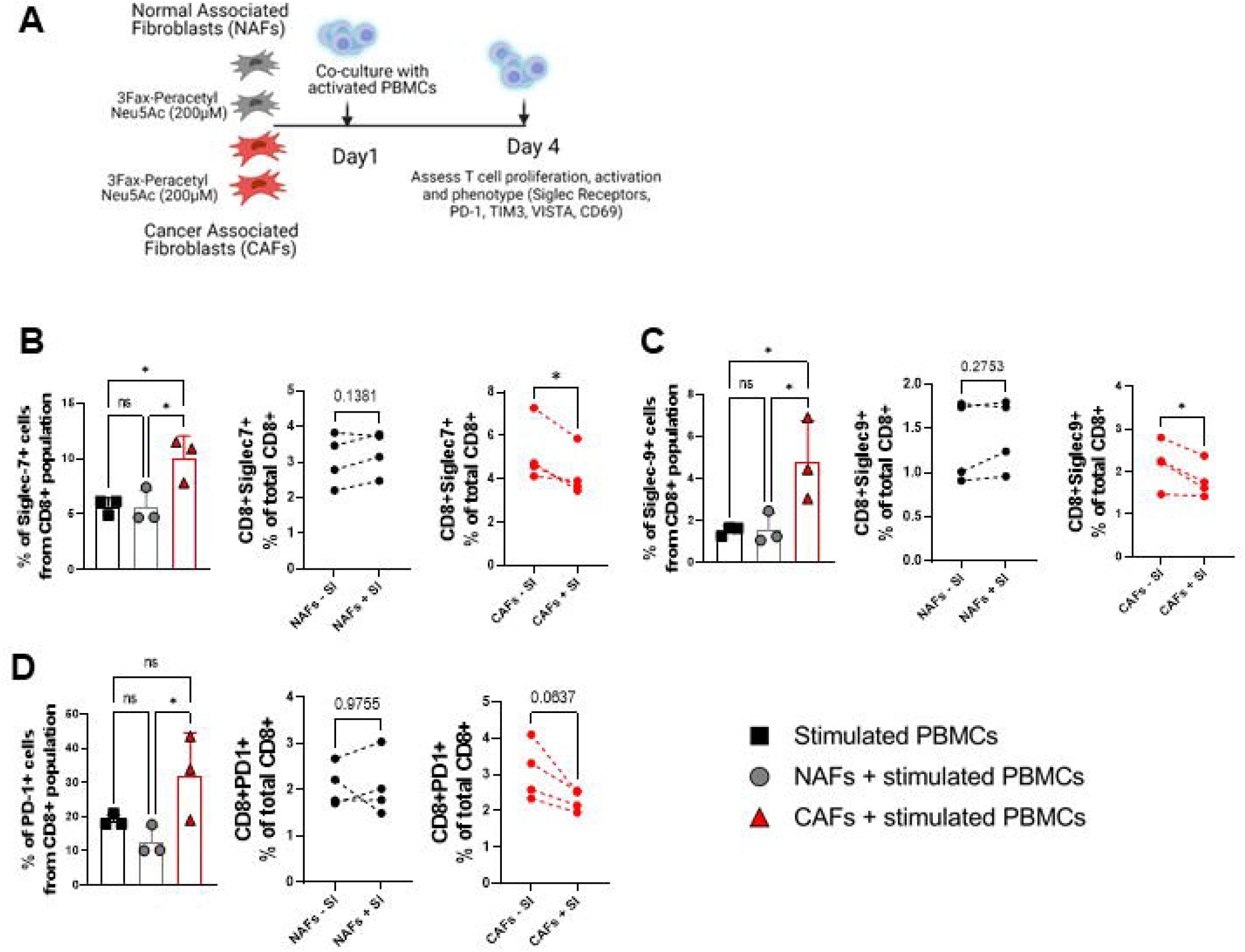
CAFs induce a sialylation dependent exhausted phenotype in CD8+ T cells. **A** NAFs and CAFs (pre-treated +/- 3Fax-PeracetylNeu5Ac (S))) were co-cultured with stimulated PBMCs **B** Frequency (%) of CD8+Siglec-7+ T cells after co-culture with NAFs or CAFs (left); Frequency (%) of CD8+Siglec-7+ T cells after co-culture with NAFs pre-treated or not with SI (middle); Frequency (%) of CD8+Siglec-7+ T cells after co-culture with CAFs pre-treated or not with SI (right). **C** Frequency (%) of Siglec-9-expressing CD8+ T cells after co-culture with NAFs or CAFs (left); Frequency (%) of CD8+Siglec-9+ T cells after co-culture with NAFs pre-treated or not with SI (middle); Frequency (%) of CD8+Siglec-9+ T cells after co-culture with CAFs pre-treated or not with SI (right) **D** Frequency (%) of CD8+PD-1+ T cells after co-culture with NAFs or CAFs (left); Frequency (%) of CD8+PD-1+ T cells after co-culture with NAFs pre-treated or not with SI (middle); Frequency (%) of CD8+PD-1+ T cells after co-culture with CAFs pre-treated or not with SI (right). Data are mean□±□SD; \**p*□<□0.05 using **(A, B and C (left))** one-way ANOVA with a Tukey post hoc test and **(B, C, D)** a ratio paired t test. *n* = 3-4 biological replicates.

## Discussion

Sialylation of tumour cells is known to induce immune suppression. By engaging inhibitory Siglec receptors and others, tumour-derived sialic acids disable major killing mechanisms of immune effector cells, trigger production of immune suppressive cytokines and dampen activation of antigen-presenting cells and subsequent induction of anti-tumour immune responses [22]. Hypersialylated tumour cells have been shown to inhibit NK activation [25, 28, 51] and recent growing data support an important role for sialylation and siglecs in polarizing macrophages from an inflammatory anti-cancer M1 to a pro-tumorigenic, immune-suppressive M2 phenotype [52, 53]. In addition, the modification of antigens with sialic acids has been shown to regulate the generation of antigen-specific Tregs via dendritic cells (DCs), leading to tolerance and inhibition of proliferation of effector T cells [54]. Targeting sialic acid and PD-1 blockade with combined checkpoint blockade proved to be synergistic, indicating immunotherapeutic potential. These observations highlight the implications of sialic acid in tumours and potential for exploiting sialic acid-siglec interactions to advance cancer immunotherapies.

The role of stromal cells, including CAFs in dictating anti-tumour immunity has been highlighted in recent studies [9, 15, 17]. However, the role of hypersialylation of stromal cells is unknown. In stromal dense tumour microenvironments, including CRC and MM, MSC/CAFs can prevent immune cell infiltration, activation and function, which facilitates tumour progression. Stromal cells modulate the immune response through the expression of immunomodulatory ligands, such as PD-L1, Fas ligand and the secretion of cytokines/chemokines [9, 12, 15, 17]. The observation that T cells are associated with stromal areas within tumours suggests that the identification of tumour stromal specific immunomodulatory mechanisms could reveal new therapeutic approaches. We demonstrate for the first time that sialic-acid-siglec signalling axis in stromal cells inhibits T cell proliferation, activation and phenotype. Importantly this effect is tumour stromal cell specific and is the first time that sialylation along with expression of functional siglec ligands have been implicated in stromal cell mediated immune modulation in cancer. Based on this data we propose that the sialylation profile of stromal cells is an important mechanism by which MSC/CAFs modulate T cell exhaustion. T cell exhaustion is associated with poor prognosis in cancer [55] and targeting CAF sialylation may represent an important therapeutic strategy to overcome this challenging feature. In addition to targeting sialylation to reverse stromal cell mediated immune suppression, it is also conceivable that this approach may also alter immune cell adhesion and trafficking in the stroma and facilitate infiltration of T cells into tumour cell regions [56]. It has recently been shown that when compared to peripheral blood lymphocytes, tumor-infiltrating lymphocytes (TILs) are more likely to express the Siglec-9 receptor as well as other checkpoint receptors such as PD-1, LAG3 and TIM3 [57]. Our findings that targeting stromal cell sialylation can reduce the frequency of Siglec 7 and 9, as well as PD-1, LAG-3 and TIM-3 expressing CD8 T cells represents a distinct advantage over targeting individual pathways of T cell exhaustion. Future studies will investigate the effects of targeting stromal cell sialyation/siglec ligands alongside checkpoint inhibitors for optimal T cell activation.

The demonstration that stromal cells express Siglec 7 and 9 ligands also has implications for a more targeted approach to disrupt Siglec/Siglec ligand stromal cell interactions. Siglec-9 ligand was upregulated on MSC/CAFs compared to their associated normal stromal cells along with upregulation of Siglec-9 on both CD4+ and CD8+ T-cells exposed to these CAFs, indicating the potential for CAFs to induce an immune suppressive effect. The significant reduction of both siglec-9 ligand on CAFs and the siglec-9 receptor on CD8+ T-cells with ST inhibitor treatment could lead to enhanced T-cell activation and immune clearance. Further research to identify specific siglec ligands could offer the potential for more specific targeting approaches for tumour associated stromal cell sialylation [26].

Potential therapeutic strategies to overcome sialylation induced immune evasion by stromal cells include the use of blocking antibodies, the targeted delivery of sialidase enzyme using conjugated antibodies and the use of ST inhibitors to block the incorporation of sialic acid onto new glycan structures. The latter approach has been pioneered by Jim Paulson’s group at Scripps, who developed the cell permeable peracetylayted fluorinated ST inhibitor 3Fax Neu5Ac, used in our study [58]. 3Fax Neu5Ac is effective at blocking the action of most sialyltransferases with minimal off target effects. From a clinical utility point of view the main challenge is on target, off tumor toxicity in the kidney, necessitating either local or targeted delivery to reduce systemic exposure and risk of nephrotoxicity [59]. Bull and colleagues showed that intra-tumoral injection of this drug in a syngeneic murine melanoma model could reverse tumor cell sialylation in-vivo and enhance T cell mediated tumor immunity [22, 30]. Following intra-tumoral injection of the ST inhibitor they observed a significant increase in NK cells, CD4 and CD8 T cells along with a significant depletion of regulatory T cells and myeloid cells. This alone was able to suppress tumor growth. Conceivably, based on our data, desialylation of MSC could have contributed to these effects. Interestingly, pre-treatment with the ST inhibitor potentiated the effect of subsequent ovalbumin specific T cells, which were adoptively transferred into mice with melanoma tumors expressing chicken ovalbumin. This suggests the potential of using ST inhibitor treatment as an adjunct to adoptive CAR-T or CAR-NK cell therapy to overcome TME immunosuppression.

An alternative approach pioneered by Bertozzi and now being pursued commercially is the use of sialidase-conjugated antibodies [60]. They observed improved in-vivo control using sialidase conjugated Trastuzumab against a HER2 + expressing tumor in a syngeneic model [61]. This anti-tumor benefit was shown to be siglec dependent, since deletion of Siglec-E, the murine homologue of Siglec-7 and 9 eliminated the difference in outcome between the two trastuzumab treatment groups, only one of whom had conjugation with an active sialidase enzyme. Future clinical studies will hopefully reveal the full value of these new glyco immune checkpoint approaches. If such strategies prove clinically useful our data suggest the benefit of not only targeting hypersialylated tumour cells, but also hyeprsialylated MSC/CAFs, especially in stromal rich tumors, where they may play a major role in immune exclusion. Using sialidase-conjugated antibodies targeting stromal cell markers may represent a clinically relevant approach to targeting sialylation and consequent immunosuppressive features in stromal dense solid tumours.

In conclusion we have demonstrated that not only are stromal cells within the tumour microenvironment highly sialylated but that sialoglycans play an important role in their immunomodulatory properties, suppressing immune cell activation, which is likely to be, at least in part, due to interaction with Siglec receptors. These data suggest that strategies to reduce sialylation of MSC could have important immune activating effects in colorectal cancer, MM and other cancers malignancies and warrant further investigation.

## Acknowledgements

This study was supported by a Science Foundation Ireland Frontiers for the Future Award (19/FFP/6446) and Starting Investigator Award (15/SIRG/3456) to A.E.R and an NUI Galway College of Medicine, Nursing and Health Sciences scholarship to Hannah Egan. This work was also supported by grants from Science Foundation Ireland (12/IA/1624) to T.R. Graphics and animations generated using Biorender by Eileen Reidy MSc. All flow cytometry experiments were performed in the NUI Galway Flow Cytometry Core Facility, which is supported by funds from NUI Galway, Science Foundation Ireland, the Irish Government’s Programme for Research in Third Level Institutions, Cycle 5, and the European Regional Development Fund. The authors wish to thank the BRU (Bio-Resources Unit) technical, veterinary, and administrative staff in NUI Galway for facilitating in vivo studies and for their ongoing assistance, advice and support in animal procedures, husbandry, care and welfare.

## Author Contributions

H.E, O.T and K.L performed majority of experiments and associated data analysis, N.A.L, G.OM, M.C performed and analysed MSC experiments. K.DeV, K.V. performed experiments relating to the 5T3MM model, M.S, A.C., L.J.E, A.M.H planned and performed surgical resections and biopsies. E.M.K., K.R. and P.D performed analysis on human CRC datasets. S.H. performed immunohistochemical analysis of human CRC tumours. T.R., M.O’D. A.E.R. conceived contributed to study design, experimental planning and data interpretation.

## References

1. Pitt, J.M., et al., Targeting the tumor microenvironment: removing obstruction to anticancer immune responses and immunotherapy. Ann Oncol, 2016. 27(8): p. 1482–92.

2. Dienstmann, R., et al., Consensus molecular subtypes and the evolution of precision medicine in colorectal cancer. Nat Rev Cancer, 2017. 17(4): p. 268.

3. Galon, J., et al., Type, density, and location of immune cells within human colorectal tumors predict clinical outcome. Science, 2006. 313(5795): p. 1960–4.

4. Roelands, J., et al., Immunogenomic Classification of Colorectal Cancer and Therapeutic Implications. Int J Mol Sci, 2017. 18(10).

5. Calon, A., et al., Stromal gene expression defines poor-prognosis subtypes in colorectal cancer. Nat Genet, 2015. 47(4): p. 320–9.

6. Guinney, J., et al., The consensus molecular subtypes of colorectal cancer. Nat Med, 2015. 21(11): p. 1350–6.

7. Leblay, N., et al., Deregulation of Adaptive T Cell Immunity in Multiple Myeloma: Insights Into Mechanisms and Therapeutic Opportunities. Front Oncol, 2020. 10: p. 636.

8. Koliaraki, V., et al., Mesenchymal Cells in Colon Cancer. Gastroenterology, 2017. 152(5): p. 964–979.

9. Turley, S.J., V. Cremasco, and J.L. Astarita, Immunological hallmarks of stromal cells in the tumour microenvironment. Nat Rev Immunol, 2015. 15(11): p. 669–82.

10. O’Malley, G., et al., Mesenchymal stromal cells (MSCs) and colorectal cancer: a troublesome twosome for the anti-tumour immune response? Oncotarget, 2016. 7(37): p. 60752–60774.

11. Nishikawa, G., et al., Bone marrow-derived mesenchymal stem cells promote colorectal cancer progression via CCR5. Cell Death Dis, 2019. 10(4): p. 264.

12. Koliaraki, V., et al., The mesenchymal context in inflammation, immunity and cancer. Nat Immunol, 2020. 21(9): p. 974–982.

13. Fessler, E. and J.P. Medema, Colorectal Cancer Subtypes: Developmental Origin and Microenvironmental Regulation. Trends Cancer, 2016. 2(9): p. 505–518.

14. Garcia-Ortiz, A., et al., The Role of Tumor Microenvironment in Multiple Myeloma Development and Progression. Cancers (Basel), 2021. 13(2).

15. O’Malley, G., et al., Stromal Cell PD-L1 Inhibits CD8(+) T-cell Antitumor Immune Responses and Promotes Colon Cancer. Cancer Immunol Res, 2018. 6(11): p. 1426–1441.

16. de Jong, M.M.E., et al., The multiple myeloma microenvironment is defined by an inflammatory stromal cell landscape. Nat Immunol, 2021.

17. Lakins, M.A., et al., Cancer-associated fibroblasts induce antigen-specific deletion of CD8 (+) T Cells to protect tumour cells. Nat Commun, 2018. 9(1): p. 948.

18. Pinho, S.S. and C.A. Reis, Glycosylation in cancer: mechanisms and clinical implications. Nat Rev Cancer, 2015. 15(9): p. 540–55.

19. Reily, C., et al., Glycosylation in health and disease. Nat Rev Nephrol, 2019. 15(6): p. 346–366.

20. Rodrigues, E. and M.S. Macauley, Hypersialylation in Cancer: Modulation of Inflammation and Therapeutic Opportunities. Cancers (Basel), 2018. 10(6).

21. Bull, C., M.H. den Brok, and G.J. Adema, Sweet escape: sialic acids in tumor immune evasion. Biochim Biophys Acta, 2014. 1846(1): p. 238–46.

22. Bull, C., et al., Sialic acids sweeten a tumor’s life. Cancer Res, 2014. 74(12): p. 3199–204.

23. Crocker, P.R., J.C. Paulson, and A. Varki, Siglecs and their roles in the immune system. Nat Rev Immunol, 2007. 7(4): p. 255–66.

24. Boussiotis, V.A., P. Chatterjee, and L. Li, Biochemical signaling of PD-1 on T cells and its functional implications. Cancer J, 2014. 20(4): p. 265–71.

25. Jandus, C., et al., Interactions between Siglec-7/9 receptors and ligands influence NK cell-dependent tumor immunosurveillance. J Clin Invest, 2014. 124(4): p. 1810–20.

26. Adams, O.J., et al., Targeting sialic acid-Siglec interactions to reverse immune suppression in cancer. Glycobiology, 2018. 28(9): p. 640–647.

27. Haas, Q., et al., Siglec-9 Regulates an Effector Memory CD8(+) T-cell Subset That Congregates in the Melanoma Tumor Microenvironment. Cancer Immunol Res, 2019. 7(5): p. 707–718.

28. Hudak, J.E., S.M. Canham, and C.R. Bertozzi, Glycocalyx engineering reveals a Siglec-based mechanism for NK cell immunoevasion. Nat Chem Biol, 2014. 10(1): p. 69–75.

29. Cornelissen, L.A.M., et al., Disruption of sialic acid metabolism drives tumor growth by augmenting CD8(+) T cell apoptosis. Int J Cancer, 2019. 144(9): p. 2290–2302.

30. Bull, C., et al., Sialic Acid Blockade Suppresses Tumor Growth by Enhancing T-cell-Mediated Tumor Immunity. Cancer Res, 2018. 78(13): p. 3574–3588.

31. Lynch, K., et al., TGF-beta1-Licensed Murine MSCs Show Superior Therapeutic Efficacy in Modulating Corneal Allograft Immune Rejection In Vivo. Mol Ther, 2020. 28(9): p. 2023–2043.

32. Ryan, A.E., et al., Targeting colon cancer cell NF-kappaB promotes an anti-tumour M1-like macrophage phenotype and inhibits peritoneal metastasis. Oncogene, 2015. 34(12): p. 1563–74.

33. De Veirman, K., et al., Multiple myeloma induces Mcl-1 expression and survival of myeloid-derived suppressor cells. Oncotarget, 2015. 6(12): p. 10532–47.

34. Vanderkerken, K., et al., Organ involvement and phenotypic adhesion profile of 5T2 and 5T33 myeloma cells in the C57BL/KaLwRij mouse. Br J Cancer, 1997. 76(4): p. 451–60.

35. Bankhead, P., et al., QuPath: Open source software for digital pathology image analysis. Sci Rep, 2017. 7(1): p. 16878.

36. Nishida, N., et al., Microarray analysis of colorectal cancer stromal tissue reveals upregulation of two oncogenic miRNA clusters. Clin Cancer Res, 2012. 18(11): p. 3054–70.

37. Ferrer-Mayorga, G., et al., Vitamin D receptor expression and associated gene signature in tumour stromal fibroblasts predict clinical outcome in colorectal cancer. Gut, 2017. 66(8): p. 1449–1462.

38. Cerami, E., et al., The cBio cancer genomics portal: an open platform for exploring multidimensional cancer genomics data. Cancer Discov, 2012. 2(5): p. 401–4.

39. Gao, J., et al., Integrative analysis of complex cancer genomics and clinical profiles using the cBioPortal. Sci Signal, 2013. 6(269): p. pl1.

40. Ren, G., et al., Mesenchymal stem cell-mediated immunosuppression occurs via concerted action of chemokines and nitric oxide. Cell Stem Cell, 2008. 2(2): p. 141–50.

41. Lin, W.D., et al., Sialylation of CD55 by ST3GAL1 Facilitates Immune Evasion in Cancer. Cancer Immunol Res, 2021. 9(1): p. 113–122.

42. Wu, X., et al., Sialyltransferase ST3GAL1 promotes cell migration, invasion, and TGF-beta1-induced EMT and confers paclitaxel resistance in ovarian cancer. Cell Death Dis, 2018. 9(11): p. 1102.

43. Guerrero, P.E., et al., Knockdown of alpha2,3-Sialyltransferases Impairs Pancreatic Cancer Cell Migration, Invasion and E-selectin-Dependent Adhesion. Int J Mol Sci, 2020. 21(17).

44. Maurice, P., et al., New Insights into Molecular Organization of Human Neuraminidase-1: Transmembrane Topology and Dimerization Ability. Sci Rep, 2016. 6: p. 38363.

45. Murphy, N., et al., TNF-alpha/IL-1beta-licensed mesenchymal stromal cells promote corneal allograft survival via myeloid cell-mediated induction of Foxp3(+) regulatory T cells in the lung. FASEB J, 2019. 33(8): p. 9404–9421.

46. Allegra, A., et al., Lymphocyte Subsets and Inflammatory Cytokines of Monoclonal Gammopathy of Undetermined Significance and Multiple Myeloma. Int J Mol Sci, 2019. 20(11).

47. van de Wall, S., et al., Sialoglycans and Siglecs Can Shape the Tumor Immune Microenvironment. Trends Immunol, 2020. 41(4): p. 274–285.

48. Laubli, H. and A. Varki, Sialic acid-binding immunoglobulin-like lectins (Siglecs) detect self-associated molecular patterns to regulate immune responses. Cell Mol Life Sci, 2020. 77(4): p. 593–605.

49. Beltra, J.C., et al., Developmental Relationships of Four Exhausted CD8(+) T Cell Subsets Reveals Underlying Transcriptional and Epigenetic Landscape Control Mechanisms. Immunity, 2020. 52(5): p. 825–841 e8.

50. Wherry, E.J. and M. Kurachi, Molecular and cellular insights into T cell exhaustion. Nat Rev Immunol, 2015. 15(8): p. 486–99.

51. Hudak, J.E. and C.R. Bertozzi, Glycotherapy: new advances inspire a reemergence of glycans in medicine. Chem Biol, 2014. 21(1): p. 16–37.

52. Beatson, R., et al., The mucin MUC1 modulates the tumor immunological microenvironment through engagement of the lectin Siglec-9. Nat Immunol, 2016. 17(11): p. 1273–1281.

53. Beatson, R., et al., Cancer-associated hypersialylated MUC1 drives the differentiation of human monocytes into macrophages with a pathogenic phenotype. Commun Biol, 2020. 3(1): p. 644.

54. Perdicchio, M., et al., Sialic acid-modified antigens impose tolerance via inhibition of T-cell proliferation and de novo induction of regulatory T cells. Proc Natl Acad Sci U S A, 2016. 113(12): p. 3329–34.

55. Liang, R., et al., TIGIT promotes CD8(+)T cells exhaustion and predicts poor prognosis of colorectal cancer. Cancer Immunol Immunother, 2021.

56. Laubli, H. and L. Borsig, Altered Cell Adhesion and Glycosylation Promote Cancer Immune Suppression and Metastasis. Front Immunol, 2019. 10: p. 2120.

57. Stanczak, M.A., et al., Self-associated molecular patterns mediate cancer immune evasion by engaging Siglecs on T cells. J Clin Invest, 2018. 128(11): p. 4912–4923.

58. Rillahan, C.D., et al., Global metabolic inhibitors of sialyl-and fucosyltransferases remodel the glycome. Nat Chem Biol, 2012. 8(7): p. 661–8.

59. Macauley, M.S., et al., Systemic blockade of sialylation in mice with a global inhibitor of sialyltransferases. J Biol Chem, 2014. 289(51): p. 35149–58.

60. Xiao, H., et al., Precision glycocalyx editing as a strategy for cancer immunotherapy. Proc Natl Acad Sci U S A, 2016. 113(37): p. 10304–9.

61. Gray, M.A., et al., Targeted glycan degradation potentiates the anticancer immune response in vivo. Nat Chem Biol, 2020. 16(12): p. 1376–1384.

