## Supplemental files for "Stromal cell sialylation suppresses T cells in inflammatory tumour microenvironments: A new tumour stromal cell immune checkpoint?"

**Supplemental Figure 1**. **Treatment with sialyltransferase inhibitor does not alter expression of mouse MSC characterisation markers**

Representative flow cytometry histograms and bar graphs showing median fluorescence intensity for the cell surface expression of the MSC characterisation markers CD44, CD73 and CD105 on $MSC^{Control}$ and $MSC^{Control + SI}$. Data are mean + SD. $n$ = 3 replicates.

**Supplemental Figure 2. Optimization of sialyltransferase inhibitor dosage and MSC treatment regime**

**(A)** Comparison of α2,6-linked sialic acid expression and **(B)** α2,3-linked sialic acid expression on $MSC^{Control}$, $MSC^{TCS}$ and $MSC^{iTCS}$ with or without treatment with a single 200µM dose, single 400µM dose or two doses of 200µM (72h apart) of sialyltransferase inhibitor. Data are mean ± SD; $^*p < 0.05$, $^{**}p < 0.01$ and $^{***}p < 0.001$ using one-way ANOVA. $n$ = 3 replicates.

**Supplemental Figure 3. MSC size, granularity and viability is unaffected by sialyltransferase treatment**

Flow cytometric analysis of **(A)** viability, **(B)** granularity and **(C)** size of $MSC^{Control}$, $MSC^{TCS}$ and $MSC^{iTCS}$ with or without sialyltransferase inhibitor treatment. **(D)** Microscopic assessment of $MSC^{Control}$, $MSC^{TCS}$ and $MSC^{iTCS}$ morphology with or without sialyltransferase inhibitor treatment. Data are mean ± SD. $n$ = 3 replicates.

**Supplemental Figure 4. Increased α2,6-linked sialic acid expression observed on both MM patient-derived MSCs and MM cell line-conditioned MSCs**

MFI of MAL II (α2,3-linked sialic acid) and SNA I (α2,6-linked sialic acid) expression on **(A)** $MSC^{Control}$, RPMI 8226 $MSC^{TCS}$ and RPMI 8226 $MSC^{iTCS}$ and **(B)** $MSC^{Control}$, MM1S $MSC^{TCS}$ and MM1S $MSC^{iTCS}$. Representative histograms of SNA I and MAL II are also shown. **(C)** MFI of MAL II and SNA I on multiple myeloma patient-derived MSCs compared to healthy bone marrow-derived MSCs. Representative histograms of SNA I and MAL II are also shown. Data are mean ± SD; $^*p < 0.05$ using **(A** and **B)** one-way ANOVA with a Tukey post hoc test and (**D**) unpaired $t$-test. $n$ = **(A** and **B)** 1-3 biological replicates and $n$ = **(C)** 4 biological replicates.

**Supplemental Figure 5. Cell surface characterisation of CRC patient-derived NAFs and CAFs**

Scatter plot bar graphs showing MFI of the stromal markers CD73 (A), CD44 (B), CD105 (C), PDGFR- α (D), PDGFR-β (E), Podoplanin (F), and HLA-DR (G) (inset: representative histograms). Data are mean ± SD; $^*p < 0.05$ using a ratio paired t test. $n$ = 3 biological replicates.

**Supplemental Figure 6. Siglec-9 ligand expression is abrogated following SI treatment**

MFI of Siglec-9 Fc chimera expression on CAFs before and after treatment with a sialyltransferase inhibitor. Data are mean ± SD; $^*p < 0.05$ using a paired t test. $n$ = 4 biological replicates.

**Supplemental Figure 7. CAFs suppress CD4+ T cell proliferation but do not induce an exhausted CD4+ T cell phenotype**

**(A)** CD4+ T cell proliferation following co-culture with NAFs or CAFs. Representative histograms depicting proliferation profiles of CD4+ T cells after co-culture with NAFs or CAFs are also shown. **(B)** Frequency (%) of CD8+CD69+ T cells after co-culture with NAFs or CAFs. **(C)** Frequency (%) of CD4+CD69+Tim-3+ T cells after co-culture with NAFs or CAFs.

Frequency (%) of **(D)** CD4+PD-1+, **(E)** CD4+PD-1+Tim-3+ and **(F)** CD4+VISTA+ T cells after co-culture with NAFs or CAFs. **G(i)** Frequency (%) of Siglec-9-expressing CD4+ T cells after co-culture with NAFs or CAFs. **G(ii)** Frequency (%) of CD4+Siglec-9+ T cells after co-culture with NAFs pre-treated or not with SI. **G(iii)** Frequency (%) of CD4+Siglec-9+ T cells after co-culture with CAFs pre-treated or not with SI. Data are mean ± SD; *$p < 0.05$, **$p < 0.01$, ***$p < 0.001$ and ****$p < 0.0001$ using **(A**, **B, C, D, E, F** and **Gi)** one-way ANOVA with a Tukey post hoc test and **(Gii** and **Giii)** a ratio paired t test. $n$ = 3-4 biological replicates.

### Supplemental Figure 1

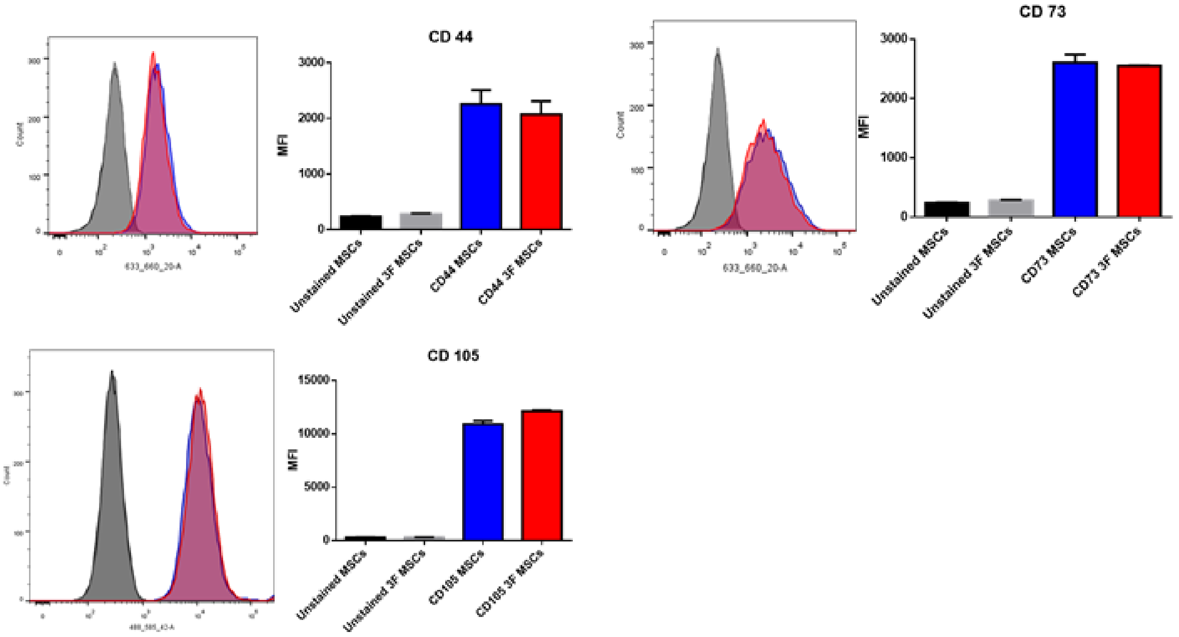

#### Supplemental Figure 2

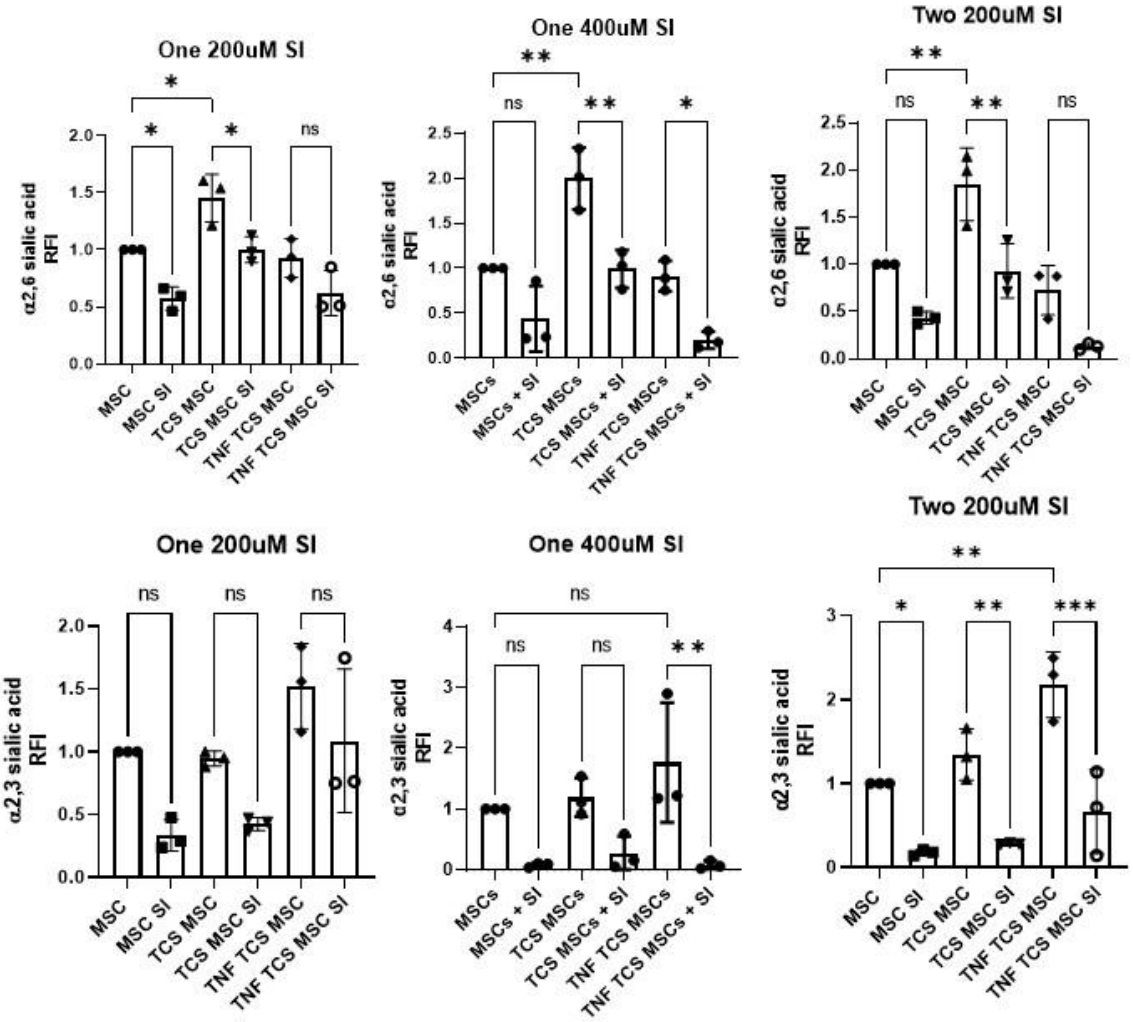

#### Supplemental Figure 3

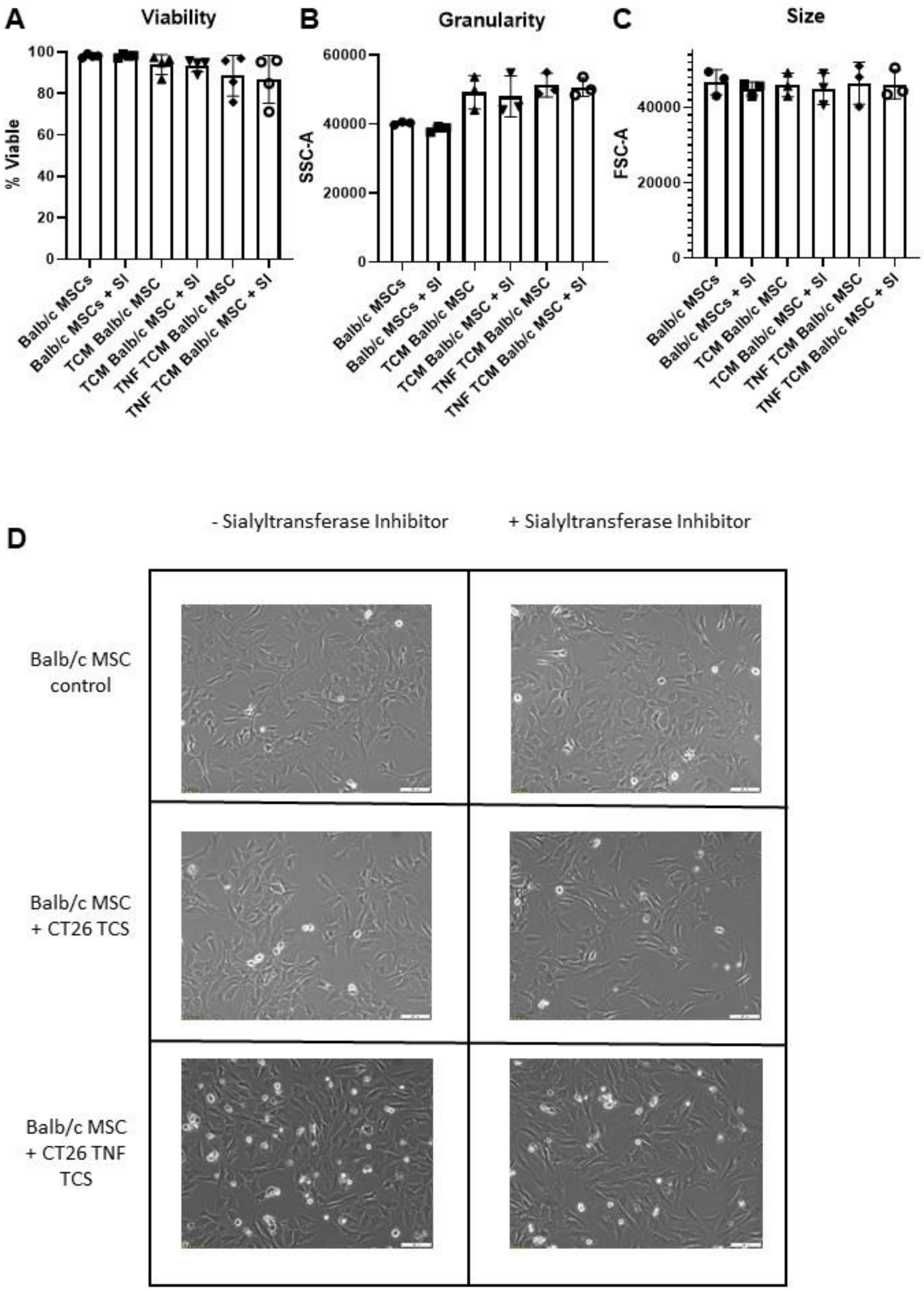

#### Supplemental Figure 4

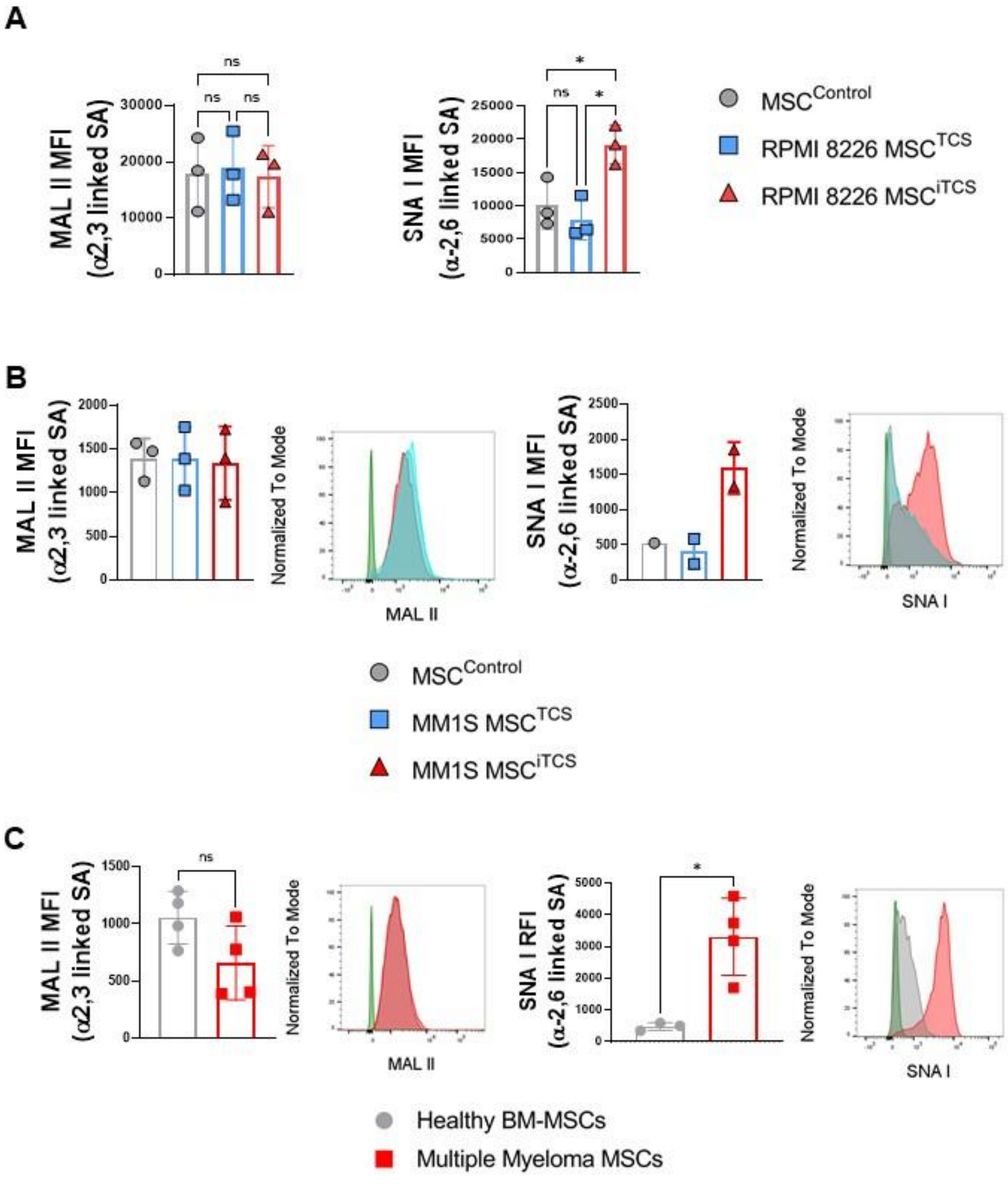

#### Supplemental Figure 5

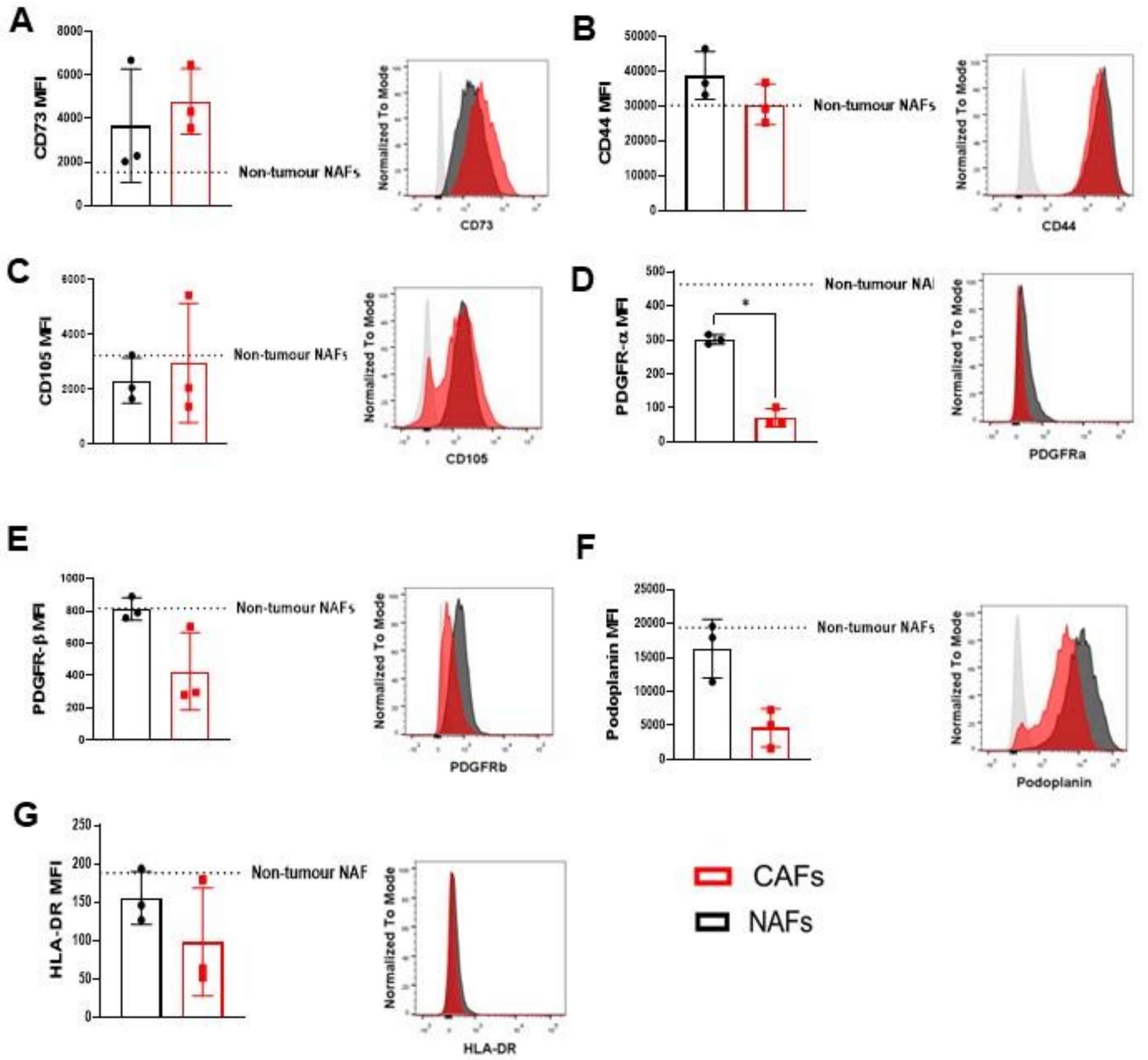

### Supplemental Figure 6

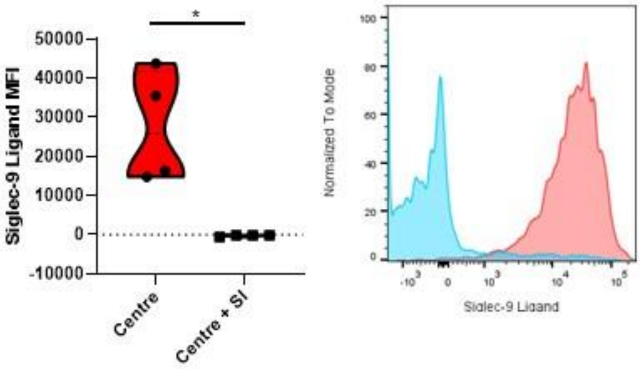

### Supplemental Figure 7

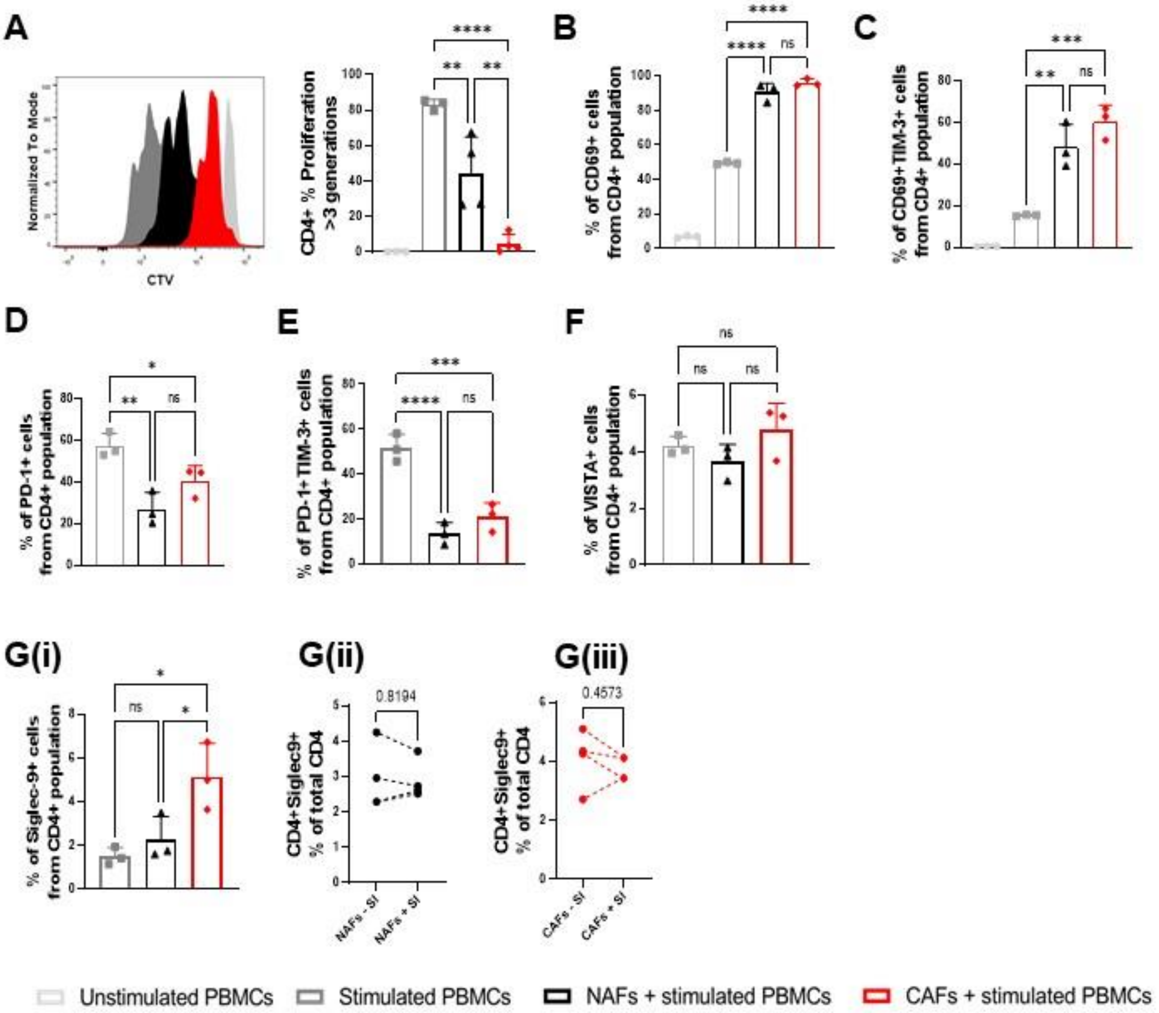
